# Comparative genomics of global *optrA*-carrying *Enterococcus faecalis* uncovers common genetic features and a chromosomal hotspot for *optrA* acquisition

**DOI:** 10.1101/785295

**Authors:** Ana R. Freitas, Ana P. Tedim, Carla Novais, Luísa Peixe

## Abstract

Linezolid-resistant *Enterococcus faecalis* (LREfs) carrying *optrA* are increasingly reported globally from multiple sources, but we still lack a comprehensive analysis of human and animal *optrA*-LREfs strains. We investigated the phylogenetic structure, genetic content [antimicrobial resistance (AMR), virulence, prophages, plasmidome] and *optrA*-containing platforms of 28 publicly available *optrA*-positive *E. faecalis* genomes from different hosts in 7 countries. In the genome-level analysis, *in house* databases with 57 virulence and 391 plasmid replication genes were tested for the first time. Our analysis showed a diversity of clones and adaptive gene sequences related to a wide range of genera, mainly but not exclusive from *Firmicutes*. The content in AMR and virulence genes was highly identical in contrast to the diversity of phages and plasmids observed. Epidemiologically unrelated clones (ST476-like and ST21-like) obtained from human clinical and animal hosts in different continents over 5 years (2012-2017) were phylogenetically related (3-122 SNPs difference). They also exhibited identical AMR and virulence profiles, highlighting a global spread of *optrA*-positive strains with relevant adaptive traits in livestock and that they might originate from an animal reservoir. *optrA* was located on the chromosome within a Tn*6674*-like element (n=9) or on medium-size plasmids (30-60 kb; n=14) belonging to main plasmid families (RepA_N/Inc18/Rep_3). In most cases, the immediate gene vicinity of *optrA* was identical in chromosomal (Tn*6674*) and plasmid (*impB-fexA-optrA*) backbones. Tn*6674* was always inserted in the same Δ*radC* integration site and embedded in a 32 kb chromosomal platform common to diverse strains from different origins (patients, healthy humans, and animals) in Europe, Africa, and Asia during 2012-2018. This platform is conserved among hundreds of *E. faecalis* genomes and we here propose a conserved chromosomal hotspot for *optrA* integration. The finding of *optrA* in strains sharing identical adaptive features and genetic backgrounds across different hosts and countries suggest the occurrence of common and independent genetic events occurring in distant regions, and might explain the easy *de novo* generation of *optrA*-positive strains. It also anticipates a dramatic increase of *optrA* carriage and spread with a serious impact in the efficacy of linezolid for the treatment of Gram-positive infections.

## Introduction

Oxazolidinones (linezolid and tedizolid) are increasingly used for the treatment of human infections caused by relevant pathogens such as methicillin-resistant *Staphylococcus aureus* and vancomycin-resistant enterococci [1, 2]. Linezolid resistance rates remain generally low in enterococci causing infections worldwide (<1%) [1, 3], however acquired linezolid resistance genes (*cfr, optrA* and/or *poxtA*) are being increasingly reported in different enterococcal species and across different settings [2–7]. Among the three transferable linezolid resistance genes, *optrA* has been the main responsible for the recent increase in linezolid-resistant enterococci (LRE) in human isolates [8–13]. According to available studies, *optrA*-carrying LRE are globally circulating in hospitals since at least 2005 [14], while the first description in food-producing animals dates from 2008 [15].

*optrA* codifies for an ABC-F protein targeting the ribosome of Gram-positive bacteria and mediates resistance to both phenicols and oxazolidinones through ribosomal protection [16]. As phenicols are allowed in veterinary medicine to control respiratory infections, the use of florfenicol in food-producing animals might co-select resistance for oxazolidinones and promote the occurrence of an *optrA* animal reservoir [7]. *optrA*-carrying LRE strains were already described in a variety of hosts including hospitalized humans, healthy humans and different food-producing animals (poultry, pigs, cattle and ducks) worldwide, and embedded in a high diversity of genetic backgrounds in association with different chromosomal or plasmid genetic platforms [1–3, 14, 15, 17–19]. Moreover, *optrA* has been more commonly described in *Enterococcus faecalis* (Efs) than in *Enterococcus faecium*.

Even though *optrA*-carrying LRE strains pose a growing threat to public health, little has been done regarding a global comparative genomics analysis and detailed *optrA* genetic context characterization [5, 9, 20, 21]. We aimed to comprehensively characterize and compare publicly available *optrA*-carrying *E. faecalis* genomes in order to investigate about the pathogenic potential of circulating clones and *optrA* mobilization.

## Material and Methods

### Bacterial collection and comparative genomic analysis

Publicly available *optrA*-carrying *E. faecalis* genomes (n=28) in the GenBank database (last search March 2019) were retrieved together with all available epidemiological information (Table 1). Only complete, assembled and identifiable genomes were included in this analysis.

**Table 1.** Epidemiological background of the 28 *E. faecalis* genomes analysed in this study.

| Isolate | Source | Sample | Country | Year | ST | Bioproject | GenBank | Reference |
| --- | --- | --- | --- | --- | --- | --- | --- | --- |
| 12E | Retail chicken | Meat | Colombia | 2010-2011 | 59 | PRJEB15278 | - | 22 |
| 34E | Retail chicken | Meat | Colombia | 2010-2011 | 59 | PRJEB15278 | - | 22 |
| 745E | Retail chicken | Meat | Colombia | 2010-2011 | 489 | PRJEB15278 | - | 22 |
| 612T | Retail chicken | Meat | Tunisia | 2017 | 21 | PRJNA517381 | SEWV01000000 | 7 |
| 728T | Retail chicken | Meat | Tunisia | 2017 | 859 | PRJNA517381 | SEWT01000000 | 7 |
| 697T | Retail chicken | Meat | Tunisia | 2017 | 476 | PRJNA517381 | VDMY00000000 | 7 |
| N60443F | Retail cattle | Meat | USA | 2014 | 489 | PRJNA292669 | CP028724-6 | 5 |
| N48037F | Retail pig | Meat | USA | 2013 | 141 | PRJNA292669 | CP028720-3 | 5 |
| 712T | Chicken | Faeces | Tunisia | 2017 | 476 | PRJNA517381 | SEWU01000000 | 7 |
| P.En090 | Pig | Faeces | Malaysia | 2012 | 756 | PRJNA343055 | MJBX01000000 | 23 |
| Enfs51 | Pig | Faeces | Malaysia | 2012 | 754 | PRJNA342925 | MJEK01000000 | 23 |
| P.En250 | Pig | Faeces | Malaysia | 2012 | 476 | PRJNA343058 | MJBZ01000000 | 23 |
| Enfs85 | Pig | Faeces | Malaysia | 2012 | 755 | PRJNA343052 | MJBV01000000 | 23 |
| IBUN9046YE | Pig | Faeces | Colombia | 2014 | 16 | PRJNA381315 | NCAD01000000 | - |
| 12T | Wastewaters | Wastewaters | Tunisia | 2014 | 86 | PRJNA386958 | NHNF01000000 | 24 |
| P.En218 | Human (Pig Farmer) | Faecal sample | Malaysia | 2012 | 765 | PRJNA343057 | MJBY01000000 | 23 |
| Enfs94 | Human (Pig Farmer) | Faecal sample | Malaysia | 2012 | 227 | PRJNA343053 | MJBW01000000 | 23 |
| Efs 599 | Healthy human | Unknown | USA | Unknown | 16 | PRJNA81957 | ALZI01000000 | - |
| 11340 | Hospitalized patient | Ascites | China | 2013 | 476 | PRJNA478627 | QNGJ01000000 | 25 |
| 13484* | Hospitalized patient | Fester | China | 2013 | 21 | PRJNA478627 | QNGN01000000 | 25 |
| 18026 | Hospitalized patient | Ascites | China | 2014 | 476 | PRJNA478627 | QNGT01000000 | 25 |
| 27149* | Hospitalized patient | Ascites | China | 2014 | 858 | PRJNA478627 | QNHA01000000 | 25 |
| Iso1Bar | Hospitalized patient | Urine | Spain | 2016 | 585 | PRJEB21295 | - | 12 |
| Iso2Bar | Hospitalized patient | Urine | Spain | 2016 | 585 | PRJEB21295 | - | 12 |
| Iso3Bar | Hospitalized patient | Urine | Spain | 2016 | 585 | PRJEB21295 | - | 12 |
| Iso4Bar | Hospitalized patient | Urine | Spain | 2016 | 474 | PRJEB21295 | - | 12 |
| Iso5Bar | Hospitalized patient | Urine | Spain | 2016 | 474 | PRJEB21295 | - | 12 |
| 16-372 | Hospitalized patient | Urine | France | 2016 | 480 | PRJNA419534 | PHLF01000000 | 10 |

Abbreviations: ST, sequence type. \*These isolates were obtained from hospitalized patients that work as farmers.

They were screened for genes encoding antibiotic resistance (ABR), virulence and MLST using *in silico* genomic tools (ResFinder 3.0, VirulenceFinder 1.5 and MLST 1.8 tools, respectively) available at the Center for Genomic Epidemiology (CGE; http://www.genomicepidemiology.org). As the VirulenceFinder database is not complete for the *E. faecalis* species (n=22 virulence genes typical of this species), we created an *in-house* database with 35 additional virulence genes and tested it using the MyDbFinder (BLAST) tool available at CGE, making a total of 57 virulence genes. The 28 assembled genomes were also analyzed by PHASTER (PHAge Search Tool Enhanced Release) to identify the presence of prophage sequences [26]. Genomic data for 23S rDNA mutations and for ribosomal protein genes *rplC, rplD* and *rpIV* were compared to reference gene loci of the linezolid-susceptible *E. faecalis* V583 (GenBank acc. no. AE016830) using Geneious Prime BLASTN. The presence of *cfr* variants, *poxtA* and plasmid replicons was also performed using Geneious Prime BLASTN and reference *cfr/poxtA* sequences [GenBank accession numbers AM408573 (*cfr*1), AJ879565 (*cfr*2), NG_055643 (*cfr*C), PHLC01000011 (*cfr*D) and MF095097 (*poxtA*)] and an *in house* database using the 391 plasmid replication sequences described in Lanza *et al*. [27]. All 28 genomes were mapped against the *E. faecalis* V583 reference strain to infer a phylogeny based on the concatenated alignment of high-quality single nucleotide polymorphisms (SNPs) using CSI Phylogeny standard settings (CGE, https://cge.cbs.dtu.dk/services/CSIPhylogeny). The phylogenetic tree was plotted using the iTOL tool [28].

### Characterization of optrA genetic platforms

The *optrA*-containing contig of the 612T genome was annotated using Geneious Prime and local database of GenBank extracted *E. faecalis* genomes. Annotation was confirmed using BLASTN, BLASTP and ISFinder. The same strains used for comparative genomics analysis were mapped against the *optrA*-containing contig of 612T using Geneious Prime in order to determine the presence of the same *optrA* platform and the genome region where it was inserted. Vector NTI advance v11 and Easyfig v2.2.2 [29] were used for drawing *optrA* platform schemes and contig comparisons. Gene functions of the 612T *optrA*-containing contig and of the different *optrA* platforms were classified according with the database of Clusters of Orthologous Groups of proteins (COGs) using eggNOG 4.5.1 (http://eggnogdb.embl.de/#/app/home). The identification of contigs belonging to the *optrA*-containing plasmid in 728T strain and the corresponding annotation was determined by comparison with the GenBank database using BLASTN, BLASTP and ISFinder. Vector NTI advance v11 and EggNOG 4.5.1 were used to determine gene function and represent the different plasmid contigs.

## Results and discussion

### Phylogenetic relatedness of optrA-carrying genomes

Figure 1 represents the phylogenetic tree constructed with the entire 28 *optrA*-carrying *E. faecalis* genomes using as reference the genome of *E. faecalis* V583. The healthy humans included correspond to a human volunteer (Efs 599) of the NIH Human Microbiome Project [30] and two swine farmers (Enfs94/P.En218) working in pig farms from Malaysia [23]. In the four *E. faecalis* genomes from hospitalized patients in China, linezolid resistance was not associated with the consumption of linezolid and an asymptomatic gut colonization was speculated by the authors [25]. Moreover, two of them (13484/27149) were recovered from Chinese farmers [25]. Only after initial analysis we realized that the Enfs94 strain (GenBank acc. no. MJBW01000000) lacked *optrA*, but as it was included in the same research project (“*Gastrointestinal health of animal handlers with special emphasis on potential zoonotic transmission of bacterial infection*”) of other related *optrA*-carrying *E. faecalis* isolated from pigs, we maintained it for comparison [23].

**Figure 1.**
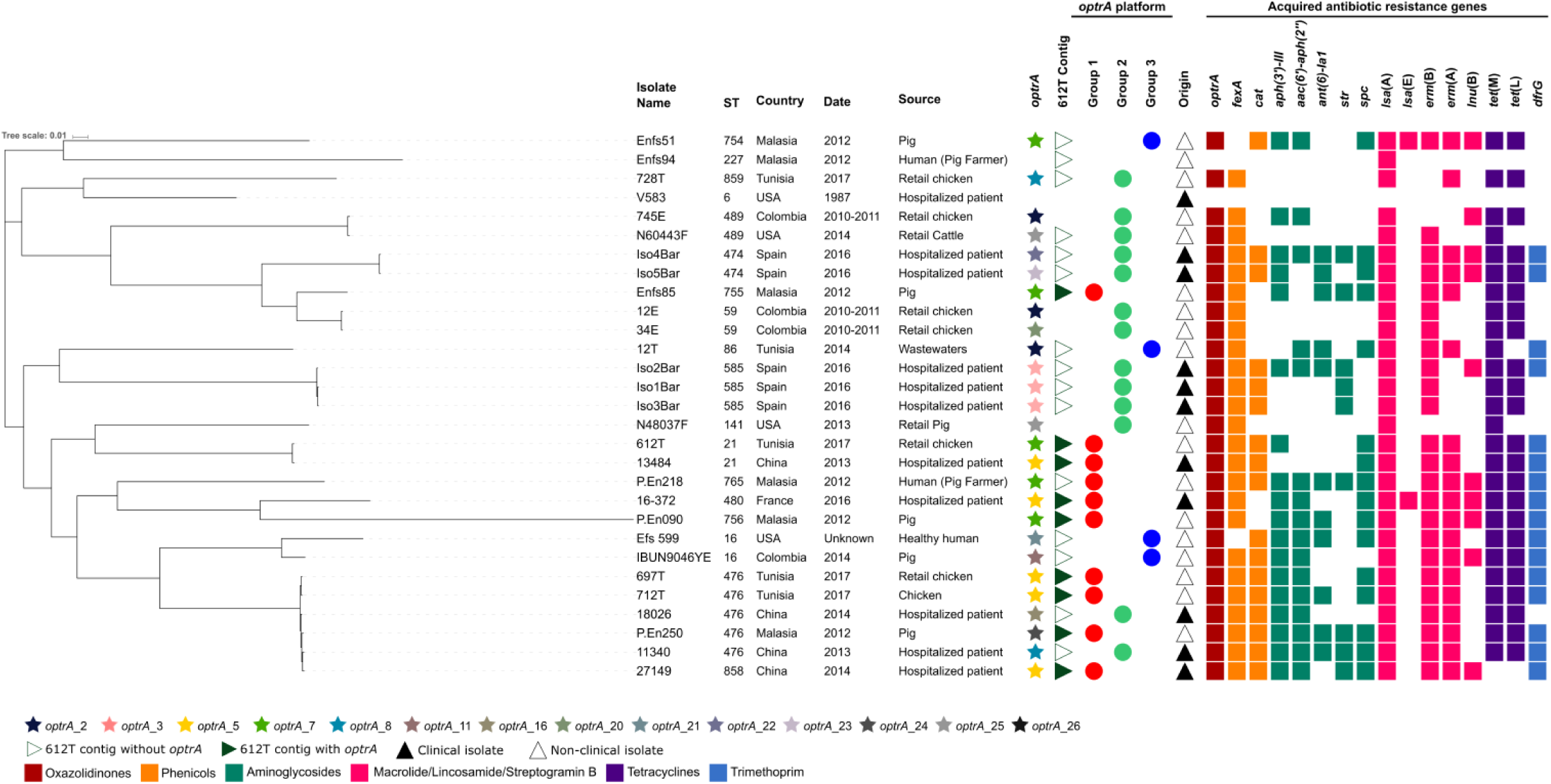
Phylogenetic tree of the *E. faecalis* strains (n=28) analysed in this study. Maximum likelihood SNP tree obtained using standard conditions of CSI Phylogeny and *E. faecalis* strains V583 as reference strain. Stars indicate the presence of the *optr*A gene. The colour of the star indicates the different *optrA* variants as indicated in the key. Triangle filled or not filled with dark-green respectively indicates the presence or absence of the chromosomal 612T contig containing *optr*A. The different *optrA* genetic platforms are represented with circles filled with red (group 1), green (group 2) and blue (group 3) as in Figure 2. Coloured cells represent the presence of acquired AMR genes with each colour indicating the correspondent family as indicated in the key (oxazolidinones in red; phenicols in yellow; aminoglycosides in green; macrolides, lincosamides or streptogramins in pink; tetracyclines in purple; and trimethoprim in blue). AMR genes present in only one isolate are not represented in the figure: *cfr* and *poxtA* (Efs599), *tet*(O) (728T) and *tet*(K) (12E). Abbreviations: ST, sequence type. White cells represent the absence of genes.

Based on SNP analysis, we observed an overall diverse population of *optrA*-positive *E. faecalis* (range 0-17.712 SNPs) with a correspondence between the sequence types identified (17 STs, 18 branches) (Figure 1, Table S1 in the Supplemental material). Strains originating from the same source or country were dispersed across the different tree branches, suggesting no significant correlation between strain origin and phylogeny as commonly observed in *E. faecalis* [31]. These results further confirm the easy promiscuity of *optrA* that has been identified in enterococci from highly variable epidemiological and genetic backgrounds [2, 32]. The exceptions corresponded to specific strains originating both temporally and locally from the same scenario namely the clinical isolates Iso1Bar/Iso2Bar/Iso3Bar (15-35 SNPs) or Iso4Bar/Iso5Bar (69 SNPs) from an hospital in Barcelona, and the chicken meat isolates 12E/34E (48 SNPs) from Colombia, with close clonal relationships as recently described [12, 22]. Still, two related clusters comprising strains from different origins and lacking evident epidemiological relationships were identified and included: i) five ST476 and one ST858 (single locus variant of ST476) isolates from chicken faeces and chicken meat (712T and 697T, respectively) in Tunisia, pigs in Malaysia (P.En250) and hospitalized patients in China (11340, 18026, 27149) differing between 3-122 SNPs; and ii) two ST21 isolates from chicken meat in Tunisia (612T) and a hospitalized patient in China (13484) differing in 113 SNPs (Table S1 in the Supplemental material). and ii) two ST21 isolates from chicken meat in Tunisia (612T) and a hospitalized patient in China (13484) differing in 113 SNPs (Table S1 in the Supplemental material).

Taking in consideration that homologous recombination events can result in numerous SNPs and genes not present in the reference strain are not taken into account (limitations of the method), these isolates could be even more closely related. In any case, the finding of the related *optrA E. faecalis* strains (ST21 and ST476) in human clinical and animal hosts from different continents over >5 years (2012-2017) was remarkable and demonstrates that they might originate from an animal reservoir and are able to easily colonize humans. An inter-hospital spread of ST476 *optrA*-carrying *E. faecalis* in different Chinese cities has been previously reported [21]. Noteworthy, most of the food animal strains included in this phylogenetic analysis (ST21, ST59, ST86, ST141, ST227, ST474, ST476, ST489, ST585) have been previously associated with *E. faecalis* from human infections in different continents, linked to linezolid resistance or not [www.pubmlst.org; [9, 12, 13, 33]].

### Comparative genomic analysis of optrA-carrying E. faecalis: antibiotic resistance, virulence, prophages and plasmid replicons

Common patterns of AMR and virulence genes were depicted among the 28 genomes, whereas that of prophages and plasmids were highly variable (Figure 1 and Figure 2). The co-location in the same contig of several of these adaptive genes encoding for AMR, virulence and plasmids (data not shown) in most genomes suggests that they may transfer easily together between different isolates.

**Figure 2.**
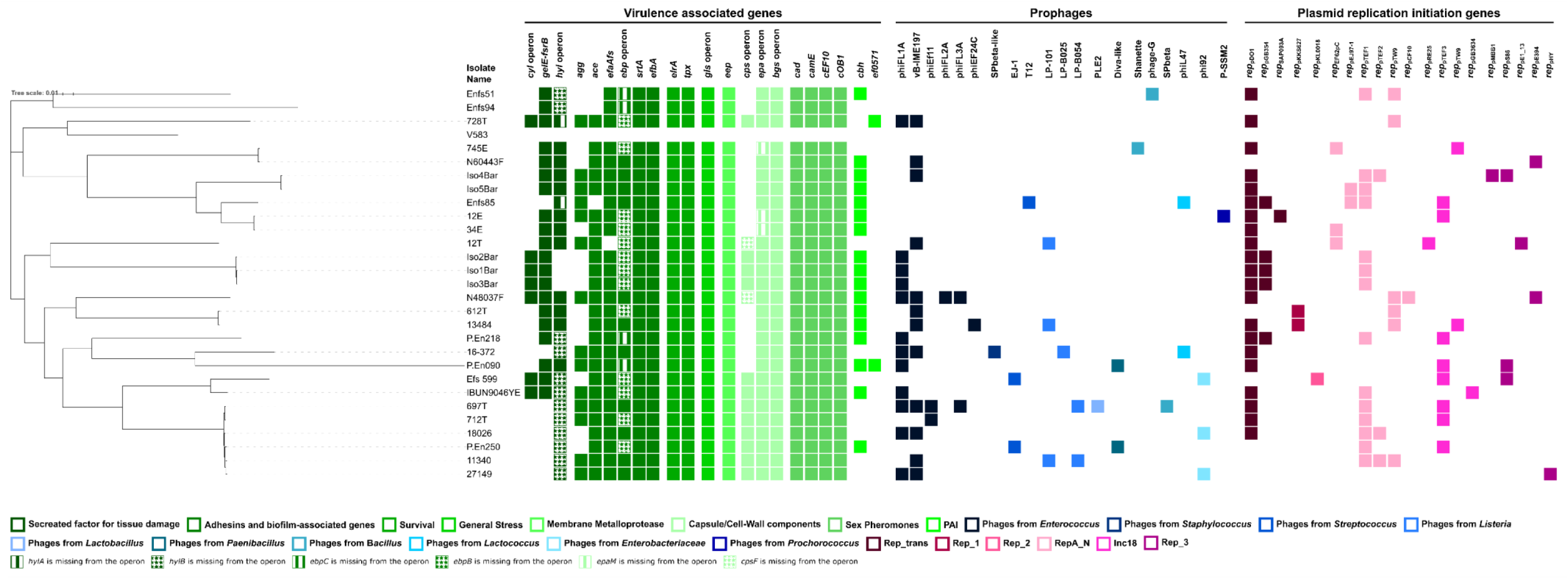
Profiles of virulence, prophages and replication initiation genes present in the *E. faecalis* (n=28) strains analysed in this study. Maximum likelihood SNP tree obtained using standard conditions of CSI Phylogeny and *E. faecalis* strains V583 as reference strain. The presence of the virulence genes, prophages and plasmid replication initiation genes is indicated by coloured squares as indicated in the key (virulence genes in green; prophages in blue; replication initiation genes in pink) and white cells indicate the absence of genes.

#### a) Antibiotic Resistance

The overall acquired ABR gene profiles identified are the same as that commonly identified in *optrA*-carrying *E. faecalis* strains from non-clinical sources in different regions [15, 18], with no significant differences between clinical (8-15 genes, 11 in average) and non-clinical (1-15 genes, 10 in average) isolates (Figure 1). They included acquired genes putatively conferring resistance to oxazolidinones, phenicols, aminoglycosides, macrolides, lincosamides or streptogramins, tetracyclines, and trimethoprim, and varying in their homology against reference strains (Table S2 in the Supplemental material). Chromosomal point mutations putatively conferring resistance to linezolid (in 23S rRNA and L3/L4/L22 ribosomal proteins) and ciprofloxacin [*gyrA* (S83I, E87G) and/or *parC* (S80I)] were also identified in most strains. Regarding linezolid resistance mutations, only synonymous ones were identified in *rpl*C/L3 and *rpl*V/L22 ribosomal proteins while in the case of the 23S rRNA some strains (n=10/28) exhibited non-synonymous mutations not previously described (Table S2 in the Supplemental material). Similarly to previous studies, all *E. faecalis* strains exhibited an amino acid substitution at position 101 of the *rpl*D/L4 protein, which has been linked to both linezolid-resistant and susceptible isolates [7, 9, 24]. Interestingly, the *optrA*-negative Enfs94 is described as linezolid resistant (MIC not available) according with Tan *et al*. [23], and so its 23S rRNA (G388A/D130N) and *rpl*D/L4 protein (T301C/F101L) non-synonymous mutations might contribute to linezolid resistance expression as this strain does not have any other known linezolid resistance mechanism. Curiously, strains from healthy humans presented very different ABR profiles: ST227 strain from a pig farmer (Enfs94) only carried *lsa*(A) and the ST16 USA human (Efs 599) carried the oxazolidinones resistance genes *poxtA* and *cfr* in addition to *optrA*. This last isolate Efs 599 (GenBank acc. no. EJU90935) carrying *optrA, poxtA* and *cfr* was obtained from a sample characterized as obtained from a “normal” volunteer of the NIH Human Microbiome Project, though no further information is available.

The two related clusters of ST476- and ST21-like strains from foodborne animals and hospitalized patients exhibited similar profiles of both acquired ABR genes and point mutations, with genes conferring resistance to aminoglycosides being the more variably present among the different strains. Enterococci displaying the referred ciprofloxacin point mutations in *gyrA/parC* have been commonly identified in samples obtained from animals and foodstuffs, but also from human infections [34, 35]. Interestingly, some of the strains clustering into the same branch in the phylogenetic tree shared similar combinations of *gyrA/parC* mutations but the same was not observed for linezolid chromosomal mutations or *optrA* variants which greatly varied even between closely related strains (Figure 1, Table S2 in the Supplemental material).

#### b) Virulence gene profiles

We herein firstly provide an *in silico* analysis of an extended number of putative virulence genes (n=57), as the few available studies analysing virulence generally include a small set of 22 genes available in the VirulenceFinder database [12, 22, 24]. The genomes here analysed contained a quite high number of virulence genes (37-54, 44 in average), though with variable gene identity in comparison to reference strains and some in truncated or increased size versions (Table S3 in the Supplemental material). Virulence profiles were generally identical beweteen clinical (38-50 genes) and non-clinical (37-54 genes) isolates (Figure 2). Both the related clusters comprising ST476-like and ST21-like strains exhibited identical virulence gene profiles including 46-47 and 40-41 genes, respectively.

Thirty-one out of the 55 virulence genes detected were present in all strains and they were related with adhesion, survival, general stress, cell surface behaviour, capsule production and sex pheromones. Interestingly, all strains carried four major virulence genes or operons relevant for *E. faecalis* endocarditis, including: *bgsA/bgsB* (glucosyltransferases with a key role in the production of the major cell wall glycolipids, biofilm production and presumably in endocarditis) [36], *gls24/glsB* (stress genes important for survival *in vivo*, and *gls24* in particular to bile salts resistance and hypothetically for virulence in peritonitis and endocarditis) [37], *efbA* (a PavA-like fibronectin adhesin previously shown to be important in experimental urinary tract infection and endocarditis) [38], and *eep* (a membrane metalloprotease crucial in biofilm formation and required for endocarditis) [39], all exhibiting nucleotide identities varying between 97% and 100% (Table S3 in the Supplemental material). Except for *epaM* missing in three strains, all genes comprising the *epa* locus (*epaA-epaR*) were found in all strains. *epa* locus is required for the biosynthesis of an antigenic rhamnopolysaccharide that plays a role in *E. faecalis* urinary tract infections and peritonitis, in intestinal colonization and in protecting against human host defenses [40]. Teng *et al*. have also described that *epa* genes are widespread among *E. faecalis* despite showing differences in sequence and cluster composition [41], but the impact of such differences on their expression remains unknown. Survival [*elrA* (leucine-rich protein A associated with macrophage persistence), *tpx* (thiol peroxidase for oxidative stress resistance)] and sex pheromone-associated (*cad, camE, cCF10, coB1*) genes were also common to all strains.

Among the six virulence gene operons identified (*hylA/hylB, ebpA/ebpB/ebpC, gls24/glsB, cpsC-cpsK, epaA-epaR* and *bgsA/bgsB*), only the *bgsA/bgsB* was complete in all strains. The six *cps* genes essential for capsule biosynthesis (*cpsC, cpsD, cpsE, cpsG, cpsI*, and *cpsK*) were identified in 14/28 strains obtained from human, environmental and animal (including the ST476 cluster) sources. The identity of these genes varied from those carried by the V583 strain (97-100% in nt), and some corresponded to pseudogenes (*cpsF* and *cpsG*; Table S3 in the Supplemental material). Non-encapsulated strains have been linked to *E. faecalis* endocarditis [42].

Although it’s hard to infer by genomics the true pathogenic potential of the 28 strains, they are undeniably enriched in diverse virulence genes playing key roles in *E. faecalis* pathogenicity and the profiles and sequences of virulence genes were generally similar between strains obtained from foodstuffs/livestock and those causing human infections (Table S3 in the Supplemental material). Moreover, specific virulence (*ace, gelE, agg, cyl, elrA*) and ABR genes (including those harboured by the genomes of our study, e.g. encoding resistance to tetracyclines, aminoglycosides, chloramphenicol, trimethoprim and MLSB) here identified have been positively associated with *E. faecalis* from human infections and belonging to dominant hospital lineages [43, 44].

#### c) Prophages

Prophages-associated sequences are commonly found in clinical strains such as the well-known V583 harbouring seven prophage-like elements, of which only two correspond to intact ones [45]. They are also commonly found in human and animal sources but the identification and distribution of phages is scarcely known in well-characterized *E. faecalis* populations [46].

Even though the overall content in phage-associated sequences of our 28 genomes was lower than that of the V583 strain, a remarkable variety of prophages was identified (20 types) and they were present in most strains (n=26/28; 1-7 prophages) with no obvious relationship with isolation source or phylogeny (Figure 2, Table S4 in the Supplemental material). Most strains (n=18/28, 64%) contained common enterococcal prophages (phiL1A and vB_IME197), but prophages associated with other bacterial genera from *Firmicutes* and in a lesser extent to *Proteobacteria* (enterobacteria) and marine *Cyanobacteria* (*Prochlorococcus*) were also identified. *Prochlorococcus* are one of the most abundant photosynthetic organisms and major producers of oxygen in plankton communities, where *E. faecalis* can survive and grow in high amounts [47].

Intact prophages were detected in 16 out of 28 strains from clinical and non-clinical sources. They putatively correspond to the *E. faecalis* vB_EfaS_IME197 (2 chicken and one wastewater strains), the *E. faecalis* phiFL1A (6 clinical, 2 pig and one human strains) and phiFL3A (one chicken strain), the *E. faecalis* phiEf11 (two chicken strain), and the *Listeria* phages LPB025 (one clinical strain) and LP-101(2 clinical and one wastewater strains). All of them are classified as temperate *Siphoviridae* bacteriophages originally obtained from variable sources (humans, farms, hospital sewage) [48–50]. The phiEf11 and LP-101 phages are not present in V583 in contrast to phiFL1A and vB_EfaS_IME197, which corresponded to the predominant prophages in the collection analysed and are also widespread in *E. faecalis* from variable sources [31]. Both phiFL1A and IME197 were originally described in clinical settings in UK and China, respectively [49, 51].

We cannot discard the possibility of an erroneous output of the “defective” prophages if these are distributed in different contigs, but the finding of several intact prophages might indicate recent integrations. In any case, the variety of prophage sequences found in such a small set of genomes was remarkable and evidences the contribution of phages to the diversity of *E. faecalis* populations by acting as vectors of different gene traits during intra- and inter-species genetic exchange [46].

#### d) Plasmidome

The extraction and analysis of plasmid replicon sequences was performed according to the recent and comprehensive review of Lanza *et al*. [27]. All but the ST227 *optrA*-negative strain from the healthy farmer in Malaysia (with no rep) carried at least one rep gene (1-5 rep genes) (Figure 2, Table S5 in the Supplemental material). The strains analysed carried plasmids belonging to all the six families known to date among enterococci, either replicating by a theta mechanism (Rep_3, Inc18, RepA_N) or by rolling circle replication (RCR: Rep_trans, Rep_1 and Rep_2) [52]. When it was possible to associate *optrA* to a specific plasmid, different plasmid types were identified (Rep_3, Inc18, RepA_N).

Within rolling-circle replicating (RCR) plasmids, most strains (21 out of 28) carried one or two replicons of the Rep_trans family, namely reppDO1 from *E. faecium* (n=21), and additionally reppGB354 from *Streptococcus agalactiae* (n=5) linked to chloramphenicol resistance (*catA7*) or repSAP093A from *S. aureus* (n=1). Rep_trans plasmids are of small size and widespread among diverse enterococcal populations, being often mobilized by other ABR conjugative theta-replicating plasmids or integrated in the chromosome [27, 53]. The content in the theta-replicating plasmids was highly variable and included replicons from pheromone-responsive RepA_N plasmids, typical of *E. faecalis*, the broad-host range Inc18, and Rep_3 which includes small non-conjugative and cryptic plasmids obtained from a variety of environments but often associated with bacteriocin production. Only the pHTβ-like plasmids were not identified as they are usually found among *E. faecium* species [53]. Notably, all but two strains carried at least one rep from Inc18 or RepA_N plasmids that correspond to the conjugative plasmids usually involved in the acquisition and spread of ABR among different *Firmicutes* genera. Within RepA_N, it is recognized that the unique pheromone-responsive plasmids are highly important in the diversification and evolution of *E. faecalis* species and the finding of truncated pTEF1 rep genes in most strains might indicate events of chromosomal integration across similar IS elements with the transfer of plasmid and large genomic DNA portions as proposed by Manson *et al*. [54].

The high variability of plasmid content found in *optrA*-carrying *E. faecalis* (Table S5 in the Supplemental material) illustrates the diverse nature of MGE that can be found in *E. faecalis* in general and in particular in *optrA*-carrying strain even in those recovered from the same sample or from different samples within a short timeframe.

### optrA variants and genetic context

The *optrA* nucleotide sequences of the 28 genomes were compared against the original *optrA* gene (*optrA_1* or *optrA*WT; GenBank acc. no. KP399637). Fourteen different variants (7 new), not including the original one, were identified (Figure 1; Table S2 in the Supplemental material). As the nomenclature of *optrA* variants is not uniformed across different studies [9, 55] and available databases (CGE), we compiled the different designations and gene mutations in Table S6 (Supplemental material) to facilitate result’s interpretation. Three different *optrA* variants were identified among the four epidemiological related chicken strains from Tunisia (*optrA_5, optrA_7 and optrA_8*) which have been also identified in different *E. faecalis* clones (e.g. ST476) from humans and pigs in China [20]. *optrA* location was inferred, when possible, from the genomic sequences. This gene was either located on the chromosomally-borne Tn*6674* (n=9) or on medium-size plasmids (30-60 Kb; n=14) belonging to the plasmid families (RepA_N, Inc18 and Rep_3) usually involved in the acquisition and spread of AMR among different genera of *Firmicutes* [27]. Independently of *optrA* localization, the analysis of the *optrA* genetic platforms carried out in in the 28 strains enabled their grouping into three main groups (Figure 2). These groups, dispersed through the branches of the phylogenetic tree, showed no obvious correlation with their strain genetic background, neither with the respective *optrA* gene variant (Figure 1). They corresponded to:

i. Group I (n=10) included three chicken strains from Tunisia (1 ST21 and 2 ST476), all the pig strains from Malaysia (ST476, ST755, ST756 and ST765) and three hospital strains from France (ST480) and China (ST21 and ST858) carrying five different *optrA* variants (Figure 3). These strains shared a chromosomal segment containing *fexA-optrA* adjacent to the entire Tn*554* from *S. aureus*, varying between 11.7-12.9Kb, which is identical to the recently identified Tn*6674* [56]. Tn*6674* is a chromosomal 12.9 kb Tn*554*-related transposon obtained from the *E. faecalis* strain E1731 (GenBank acc. no. MK737778.1) that contains 7 ORFs: *tnp*ABC and resistance genes [*spc-erm(A)-fexA-optrA*] conferring resistance to spectinomycin, macrolides, phenicols and oxazolidinones, respectively; so *fexA* and *optrA* are additionally carried in Tn*6674* in comparison to Tn*554*. As illustrated in Figure 3 that includes not only some of the genomes of this study but also *optrA* platforms available in GenBank, Tn*6674* has been driving *optrA* spread amongst hospitalized patients (France, China),
ii. Group II (n=11) included chicken strains from Colombia (2 ST59 and 1 ST489) and Tunisia (ST859), cattle and pig samples from USA (ST141 and ST489), and the clinical strains from Spain (3 ST585 and 1 ST474) and China (2 ST476) carrying eight *optrA* variants. Most of them (n=7) contained a common *impB-fexA-optrA* segment that showed high similarity to the originally described *optrA*-positive pE349 plasmid (human, ST116; GenBank acc. no. KP399637) as well as other plasmids obtained from human and pig strains in China (ST27, ST59, ST480, ST585) [20]. This plasmid backbone has been also recently described in *E. faecalis* isolates from Sweden and Ireland [3]. *impB* showed the highest homology with a type VI secretion system from *E. faecium* (GenBank acc. no. ERK33158) and has been suggested to be a plasmid-associated DNA repair gene eventually playing a role in the survival under stressful environments [57]. The ST859 chicken strain (728T) additionally carried *erm*(A) in the same contig of 12.9 Kb in common to pig ST59 (p10-2-2 plasmid) and hospitalized patient ST476 strains from China [20]. *optrA* was located on plasmids ranging from ca. 30 to 60 kb in all cases [strains from Group 2 of this study (Figure 3) and those referred from China, Colombia and the USA] [5, 12, 14, 20] belonging to RepA_N, Inc18 or Rep_3 families whenever known. Some *optrA*-carrying plasmid segments (n=5) of this group were flanked by a transposase of the IS*L3* family from *Streptococcus suis* (GenBank acc. no. WP_029876135.1), a species previously recognized as a reservoir of *optrA* [32]. healthy humans (China, Malaysia) and food-producing animals (China, Malaysia, Tunisia) isolates across different continents during 2012-2018. A similar structure was also recently described in an *optrA*-carrying ST16 *E. faecalis* from a patient in Greece (sequence not available) [11]. For the isolates and corresponding studies possible to access, this genetic platform could be positively transferred by filter mating [58].

**Figure 3.**
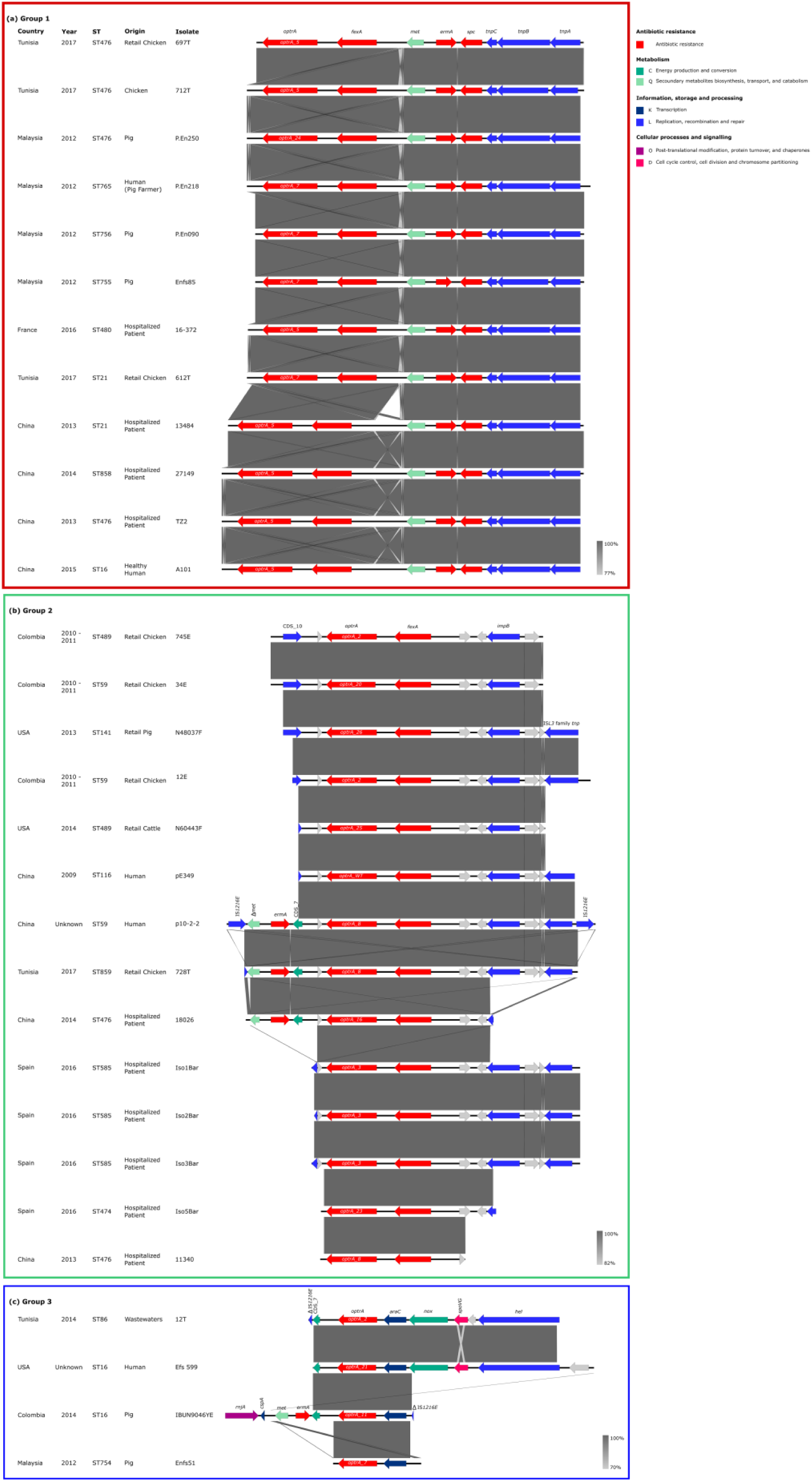
Different platforms carrying the *optr*A gene. Panel (a) group 1; (b) group 2; (c) group 3.

### Features of a novel chimeric pheromone-responsive-like optrA-carrying plasmid

We were able to presumptively identify the contigs belonging to the *optrA*-carrying plasmid of the Tunisian 728T strain from this group (Figure 4). The sum of such contigs was 39 173 bp, which is compatible with the size inferred from the S1-PFGE gel (∼40 Kb). Using EggNOG it was possible to identify the function of some of the proteins codified by the putative ORFs encoded in the plasmid, including ABR [*optrA, fexA, erm*(A), *tet*(L)], plasmid replication and stability (*repA, parA, prgN*), but not of conjugative transfer (compatible with the negative conjugation results), a toxin-antitoxin (TA) system (*mazE, mazF*), class II bacteriocin (*bacA, bacB*), cell fillamentation (*fic*), and a putative ABC-type multidrug transporter (CDS_2 and CDS_3 in the figure). The entire contig 23 of 12.971 Kb (*ermA-optrA-fexA-impB*; Figure 3; Figure 4) was identical to that from p10-2-2 plasmid obtained from a pig ST59 *E. faecalis* in China [20]. RepA had the highest homology (92% in nt) with that from the pheromone-responsive plasmids pTW9 (GenBank acc. no. NC_014726), pPD1 (GenBank acc. no. D78016) and pCF10 (GenBank acc. no. AY855841.2) previously identified in MDR *E. faecalis* strains obtained from different hosts and areas. Upstream of *repA* is *parA* and *prgN*, previously described in the pheromone-responsive pTEF2 (GenBank acc. no. AE016831) and pPD1 *E. faecalis* plasmids as putatively involved in plasmid partitioning and replication control, respectively. The region spanning from *repA* to *prgN* had the highest homology with the same *repA-parA-prgN* region of the VanB-type pMG2200 plasmid (GenBank acc. no. AB374546), still of only 73% in nucleotides, meaning this corresponds to a putative new plasmid replication region. Bacteriocin and TA gene clusters were similar to that from the pheromone-responsive pAMS1 plasmid (GenBank acc. no. EU047916) and that described in an *optrA*-positive pEF123 plasmid from chicken *E. faecalis* in China (GenBank acc. no. KX579977), respectively. The ABC transport system was described in pVEF3 (GenBank acc. no. NC_010980) and pVEF4 (GenBank acc. no. FN424376) plasmids from chicken *E. faecium* strains, though it was not associated with a reduced susceptibility to several substances tested at that time by Sletvol *et al*. [59]. Taking all together, our data suggests that this *optrA*-carrying plasmid from chicken meat corresponds to a putative novel pheromone-responsive-like plasmid containing genes of different origins including the first *optrA* pE349 plasmid described, as has been commonly observed worldwide and particularly in the non-clinical setting [2, 14, 20, 24].

**Figure 4.**
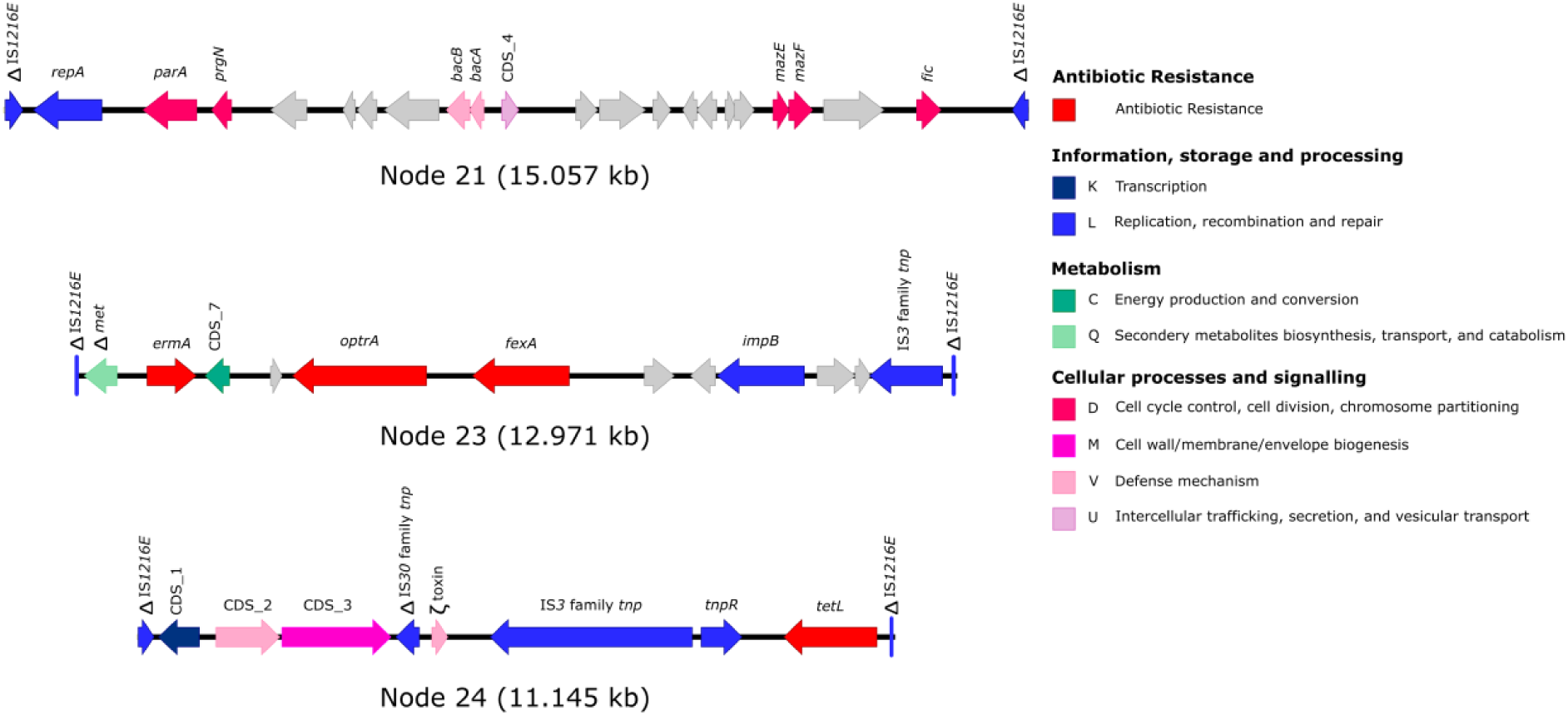
Schematic representation of plasmid of *E. faecalis* strain 728T (ST859) obtained from retail chicken in Tunisia. Coloured arrows represent genes and the arrows indicate the orientation. Grey arrows represent hypothetical proteins of unknown function; the remaining colours represent the gene function according to the COGs database classification indicated in the key. *bac*, bacteriocin; *rep*, replication; *tnp*, transposase. The presence of IS*1216* at the beginning and end of each contig made difficult to understand the order of each contig, but the similar coverage of each contig indicated that these contigs might belong to the same plasmid. A relaxase could not be identified in this plasmid, corroborating the negative conjugative assays obtained.

iii) Group III comprised four miscellaneous strains from pigs (ST16 and ST754), humans (ST16) and wastewaters (ST86). ST16 from a healthy human in USA (Efs 599) and ST86 from wastewaters (12T) in Tunisia commonly carried *optrA* adjacent to an *araC-NADH-spoVG-helicase* region highly similar to that of a blood *E. faecalis* E016 strain from China, despite the fact that in these cases *optrA* was variably located on a plasmid (ST86) or on chromosome (E016) [20, 24]. The two remaining strains (IBUN9046YE and Enfs51) only shared the *araC-optrA* segment with those aforementioned, which has been commonly described flanked by transposases and Tn*558* relics in different enterococcal species [12, 20].

#### Comparison of chromosomal *optrA*-carrying contigs

We further compared the entire chromosomal *optrA*-carrying contig (∼44 Kb) of a strain from group 1 (612T) by mapping it against the remaining genomes (Figure 5).

**Figure 5.**
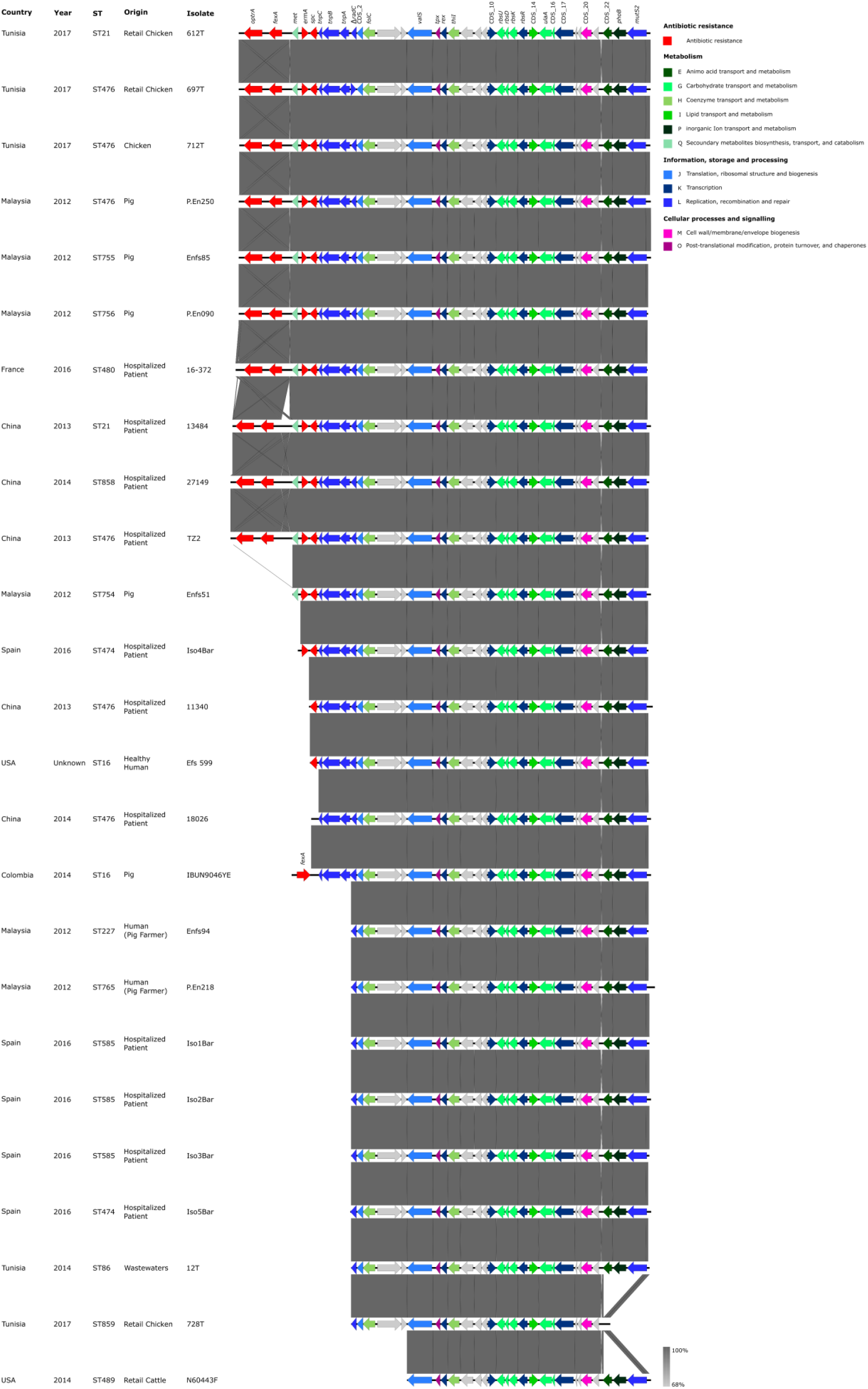
Comparison of the chromosomal 612T *optr*A-containing contig with the remaining strains. Coloured arrows represent genes and the arrows indicate the orientation. Grey arrows represent hypothetical proteins of unknown function; the remaining colours represent the gene function according to the COGs database classification indicated in the key.

In addition to the Tn*6674*-like element, this contig included genes involved in different cellular functions like oxidative stress response (*tpx*), redox-sensing transcriptional regulation (*rex*), carbohydrate transport and metabolism [*rbsUDKR* operon (ribose transport and regulation), *ulaA* (PTS system ascorbate-specific transporter)], coenzyme transport and metabolism (*folC* and *thiI* – folate and thyamine biosynthesis respectively), protein translation (*valS*, valyl-tRNA synthetase), phosphate transport and regulation (*phoB*), and DNA mismatch repair (*mutS2*). It was interesting to identify the same conserved chromosomal region (32.199 bp from Δ*radC* to *mutS2*) in all but the animal/food strains from Colombia and one from the USA (n=24/28), independently if strains carried *optrA* on this or other chromosomal contigs, on plasmids or even lacked it. *optrA* adjacent to Tn*554* or Tn*554* relics has been described either embedded on chromosomal platforms or on plasmids, in a plethora of enterococci strains from different host and regions [19, 20, 55].

As the transposons of Tn*554* family, Tn*6674* does not fall into any standard transposon category since it lacks inverted or direct terminal repeats, and it does not generate a duplication of a target sequence upon transposition. It has been suggested that Tn*554* is a prophage-like element characterized by a highly efficient site-specific integration-excision mechanism [60]. Transfer of Tn*554* between Gram-positive species, especially known between *S. aureus*, typically involves integration into another mobile element and hitch-hiking across via conjugation, transformation or transduction, so it can be found in multiple copies in the cell. Li *et al*. [56] have experimentally confirmed the ability of Tn*6674* to form circular forms and suggested its functional activity. Tn*554* has an exceptional preference for integration into a chromosomal insertion site called att*554*, at a lower frequency into secondary insertion sites (mostly in the chromosome) or more rarely on plasmids if that site is occupied or deleted, following transfer [60, 61]. The transposition of Tn*554* generally interrupts the *radC* gene encoding a putative DNA repair protein and we herein confirm the same with Tn*6674* (Δ*radC* in Figure 5) [62]. The identification of the *optrA*-carrying Tn*6674* adjacent to a common chromosomal 32 kb platform in most of the strains analysed suggests that this specific chromosomal region may be a hotspot for Tn*6674* integration and mobilization as observed for other bacterial species in which hotspots tend to be flanked by recombination and repair core genes [63]. A GenBank search of this 32 kb nucleotide sequence revealed its presence in >500 other *E. faecalis* genomes (562 out of 2701 subject genomes) from different sources (data not shown), thus demonstrating its conserved nature among this species. Remarkably, the Enfs94 strain from the healthy farmer lacking *optrA*, whom was in contact with pigs colonized with *optrA*-positive *E. faecalis*, also contained this conserved chromosomal region and an *optrA* acquisition can easily occur.

According to available studies, Tn*554* seems to transfer primarily by transducing phages independently of flanking regions [61]. Although the extent of HGT due to transduction with bacteriophages among enterococci remains largely unknown, the transfer of ABR genes between different enterococcal species mediated by phages has been already demonstrated [51, 52], as well as the effect of specific antibiotics (e.g. fluoroquinolones [45]) in prophage release and gene spread within *E. faecalis* isolates. *optrA* was not identified as integrated in a prophage in this work, but a diversity of prophages was found integrated in the chromosome of most strains including all the nine strains containing the Tn*6674* (Group 1 of Figure 3). Strikingly, phage-mediated lateral transduction was recently described as a putative new universal mechanism of transfer promoting the efficient exchange of large portions of bacterial genomes at high frequencies [64]. Considering the finding of *optrA* embedded in so many diverse genetic platforms amongst different Gram-positive genera (enterococci, staphylococci, streptococci), one can also speculate about the possible *optrA* mobilization through phage-mediated transduction. Indeed, *optrA* has already been found as integrated into a larger Φm46.1 prophage (originally from *S. pyogenes*) in *Streptococcus suis* pig isolates [32, 65]. Moreover, Wang *et al*. [66] recently demonstrated by a metagenomic approach in swine feedlot wastewater samples that phages were more likely to harbor ABC transporter family and ribosomal protection genes including *optrA*. Adding to the fact that transfer frequency of Tn*554* via transduction is higher than most plasmids [61], we could expect a tremendously fast dissemination of *optrA* among different Gram-positive species and across different hosts and settings, as seems to be occurring more expressively in the animal production scenario. It has been recently hypothesized that the virome may play a larger role in ABR transfer in non-clinical environments where phages are abundant and frequently carry ABR genes [67].

## Conclusions

This work constitutes one of the most comprehensive studies in comparing *optrA*-positive *E. faecalis* genomes from disparate origins and the first to depict extended profiles in key adaptive features including antibiotic resistance, virulence, prophages and the plasmidome. Our study evidenced the amazing variability of clones and *optrA-*genetic platforms or plasmids that can be found in a single species, and on other hand the easy possibility of finding the same clone or *optrA* genetic platform in samples from different sources and distant countries. In most cases, the immediate gene vicinity of *optrA* was identical in chromosomal and plasmid (*impB-fexA-optrA*) backbones. We herein identified a common Tn*554::fexA-optrA* chromosomal platform, the recently identified novel Tn*6674*, in different linezolid-resistant *E. faecalis* clones from livestock, meat, animal farmers and hospitalized patients across different continents, and proposed a conserved chromosomal hotspot for *optrA-fexA* integration. We also hypothesize the emergence of ST476-like clones as vehicles of *optrA* as they are increasingly identified in association with different hosts and areas, suggesting they might be particularly prone to acquire and exchange *optrA. optrA*-carrying *E. faecalis* strains from livestock/foodstuffs and human infections could be closely related and harbour indistinguishable key adaptive features, which reinforces the generalist lifestyle of this species and highlights the relevance of the animal reservoir and potential role of the food-chain in the transmission of *optrA*-positive strains. In addition, we further provide evidences that *E. faecalis* easily acquire and exchange *optrA*-carrying platforms or plasmids with other Gram-positive pathogens, besides other genetic information with a wide range of bacteria. The finding of *optrA* in strains sharing identical adaptive features and genetic backgrounds across different hosts and countries suggests the occurrence of common and independent genetic events occurring in distant regions, and might explain the easy *de novo* generation of *optrA*-positive strains. The usual association of *optrA* with Tn*554*-like or IS-like elements will certainly aid to a global and dramatic increase of *optrA* carriage and spread with a serious impact in the efficacy of linezolid for the treatment of Gram-positive infections.

## Authors’ contributions

ARF and APT designed the study. ARF and APT selected and provided characterized genomes and their corresponding epidemiological information, and analysed the genomic sequence data. ARF, APT, CN and LP analysed the data. ARF wrote the manuscript. LP provided the analysis tools and computing resources needed for the study. All authors contributed to the editing and to a critical analysis of the manuscript.

