## Supplemental material for "Comparative genomics of global *optrA*-carrying *Enterococcus faecalis* uncovers common genetic features and a chromosomal hotspot for *optrA* acquisition"

**Table S1.** Results of mapping SNPs with the CSI Phylogeny-based workflow in the 28 analysed strains.

| Isolate | | Enfs51 | Enfs94 | 728T | 745E | N60443F | Iso4Bar | Iso5Bar | Enfs85 | 12E | 34E | 12T | Iso1Bar | Iso2Bar | Iso3Bar | N48037F | 612T | 13484 | P.En218 | 16-372 | P.En090 | Efs 599 | IBUN9046YE | 697T | 712T | 18026 | P.En250 | 11340 | 27149 |
|---|---|---|---|---|---|---|---|---|---|---|---|---|---|---|---|---|---|---|---|---|---|---|---|---|---|---|---|---|---|
| | **ST** | **754** | **227** | **859** | **489** | **489** | **474** | **474** | **755** | **59** | **59** | **86** | **585** | **585** | **585** | **141** | **21** | **21** | **765** | **480** | **756** | **16** | **16** | **476** | **476** | **476** | **476** | **476** | **858** |
| **Enfs51** | **754** | 0 | 14560 | 14614 | 15127 | 15126 | 15488 | 15477 | 15260 | 15127 | 15118 | 13798 | 14208 | 14219 | 14203 | 13691 | 13780 | 13782 | 13824 | 15010 | 16362 | 14283 | 13716 | 13743 | 13742 | 13726 | 13745 | 13754 | 13758 |
| **Enfs94** | **227** | 14560 | 0 | 16020 | 16271 | 16263 | 16718 | 16704 | 16446 | 16443 | 16431 | 15696 | 15930 | 15935 | 15925 | 15660 | 15791 | 15791 | 15979 | 16921 | 17712 | 16032 | 15655 | 15761 | 15760 | 15745 | 15769 | 15780 | 15782 |
| **728T** | **859** | 14614 | 16020 | 0 | 15159 | 15158 | 15574 | 15561 | 15158 | 15040 | 15036 | 13831 | 14725 | 14732 | 14720 | 14146 | 13894 | 13889 | 14156 | 15835 | 17576 | 14684 | 13849 | 14017 | 14016 | 13998 | 14021 | 14028 | 14034 |
| **745E** | **489** | 15127 | 16271 | 15159 | 0 | 128 | 13243 | 13230 | 12482 | 11864 | 11850 | 14507 | 14875 | 14882 | 14870 | 14493 | 13868 | 13868 | 14580 | 15694 | 17299 | 14838 | 14329 | 14293 | 14292 | 14270 | 14293 | 14308 | 14309 |
| **N60443F** | **489** | 15126 | 16263 | 15158 | 128 | 0 | 13241 | 13228 | 12480 | 11864 | 11850 | 14508 | 14871 | 14878 | 14866 | 14494 | 13874 | 13874 | 14579 | 15697 | 17300 | 14843 | 14334 | 14294 | 14293 | 14271 | 14292 | 14309 | 14310 |
| **Iso4Bar** | **474** | 15488 | 16718 | 15574 | 13243 | 13241 | 0 | 69 | 6178 | 6084 | 6070 | 14877 | 14966 | 14971 | 14963 | 14814 | 14652 | 14654 | 15019 | 15723 | 17545 | 15150 | 14737 | 14924 | 14923 | 14906 | 14931 | 14938 | 14950 |
| **Iso5Bar** | **474** | 15477 | 16704 | 15561 | 13230 | 13228 | 69 | 0 | 6179 | 6081 | 6067 | 14862 | 14955 | 14960 | 14952 | 14801 | 14639 | 14643 | 15006 | 15710 | 17532 | 15137 | 14724 | 14909 | 14908 | 14891 | 14916 | 14923 | 14935 |
| **Enfs85** | **755** | 15260 | 16446 | 15158 | 12482 | 12480 | 6178 | 6179 | 0 | 3148 | 3138 | 14068 | 14581 | 14588 | 14578 | 14195 | 13790 | 13795 | 14444 | 13672 | 17359 | 13503 | 12646 | 13068 | 13067 | 13050 | 13073 | 13082 | 13086 |
| **12E** | **59** | 15127 | 16443 | 15040 | 11864 | 11864 | 6084 | 6081 | 3148 | 0 | 48 | 14208 | 14567 | 14576 | 14564 | 14082 | 13731 | 13736 | 14437 | 15537 | 17412 | 14555 | 14031 | 13919 | 13918 | 13899 | 13926 | 13933 | 13939 |
| **34E** | **59** | 15118 | 16431 | 15036 | 11850 | 11850 | 6070 | 6067 | 3138 | 48 | 0 | 14191 | 14561 | 14570 | 14558 | 14078 | 13722 | 13727 | 14432 | 15537 | 17402 | 14549 | 14023 | 13904 | 13903 | 13884 | 13911 | 13918 | 13924 |
| **12T** | **86** | 13798 | 15696 | 13831 | 14507 | 14508 | 14877 | 14862 | 14068 | 14208 | 14191 | 0 | 12830 | 12839 | 12825 | 12277 | 12408 | 12415 | 12758 | 14450 | 17418 | 13178 | 12299 | 12413 | 12412 | 12398 | 12419 | 12424 | 12436 |
| **Iso1Bar** | **585** | 14208 | 15930 | 14725 | 14875 | 14871 | 14966 | 14955 | 14581 | 14567 | 14561 | 12830 | 0 | 35 | 15 | 13020 | 12973 | 12980 | 13443 | 14669 | 17225 | 13716 | 13211 | 13220 | 13219 | 13201 | 13224 | 13239 | 13237 |
| **Iso2Bar** | **585** | 14219 | 15935 | 14732 | 14882 | 14878 | 14971 | 14960 | 14588 | 14576 | 14570 | 12839 | 35 | 0 | 30 | 13029 | 12976 | 12983 | 13450 | 14680 | 17236 | 13725 | 13220 | 13231 | 13230 | 13212 | 13233 | 13250 | 13248 |
| **Iso3Bar** | **585** | 14203 | 15925 | 14720 | 14870 | 14866 | 14963 | 14952 | 14578 | 14564 | 14558 | 12825 | 15 | 30 | 0 | 13017 | 12968 | 12975 | 13438 | 14664 | 17220 | 13713 | 13208 | 13215 | 13214 | 13196 | 13219 | 13234 | 13232 |
| **N48037F** | **141** | 13691 | 15660 | 14146 | 14493 | 14494 | 14814 | 14801 | 14195 | 14082 | 14078 | 12277 | 13020 | 13029 | 13017 | 0 | 11246 | 11254 | 12478 | 14341 | 17342 | 12348 | 11358 | 11751 | 11750 | 11731 | 11755 | 11761 | 11773 |
| **612T** | **21** | 13780 | 15791 | 13894 | 13868 | 13874 | 14652 | 14639 | 13790 | 13731 | 13722 | 12408 | 12973 | 12976 | 12968 | 11246 | 0 | 113 | 11388 | 12999 | 17464 | 12693 | 11845 | 11610 | 11609 | 11593 | 11612 | 11619 | 11627 |
| **13484** | **21** | 13782 | 15791 | 13889 | 13868 | 13874 | 14654 | 14643 | 13795 | 13736 | 13727 | 12415 | 12980 | 12983 | 12975 | 11254 | 113 | 0 | 11396 | 13001 | 17468 | 12700 | 11854 | 11613 | 11612 | 11596 | 11615 | 11624 | 11632 |
| **P.En218** | **765** | 13824 | 15979 | 14156 | 14580 | 14579 | 15019 | 15006 | 14444 | 14437 | 14432 | 12758 | 13443 | 13450 | 13438 | 12478 | 11388 | 11396 | 0 | 11316 | 17439 | 13365 | 12511 | 10610 | 10609 | 10591 | 10622 | 10623 | 10631 |
| **16-372** | **480** | 15010 | 16921 | 15835 | 15694 | 15697 | 15723 | 15710 | 13672 | 15537 | 15537 | 14450 | 14669 | 14680 | 14664 | 14341 | 12999 | 13001 | 11316 | 0 | 12780 | 9922 | 10852 | 12054 | 12053 | 12042 | 12064 | 12065 | 12067 |
| **P.En090** | **756** | 16362 | 17712 | 17576 | 17299 | 17300 | 17545 | 17532 | 17359 | 17412 | 17402 | 17418 | 17225 | 17236 | 17220 | 17342 | 17464 | 17468 | 17439 | 12780 | 0 | 15413 | 17289 | 17412 | 17411 | 17392 | 17416 | 17425 | 17424 |
| **Efs 599** | **16** | 14283 | 16032 | 14684 | 14838 | 14843 | 15150 | 15137 | 13503 | 14555 | 14549 | 13178 | 13716 | 13725 | 13713 | 12348 | 12693 | 12700 | 13365 | 9922 | 15413 | 0 | 3407 | 9901 | 9900 | 9892 | 9913 | 9918 | 9922 |
| **IBUN9046YE** | **16** | 13716 | 15655 | 13849 | 14329 | 14334 | 14737 | 14724 | 12646 | 14031 | 14023 | 12299 | 13211 | 13220 | 13208 | 11358 | 11845 | 11854 | 12511 | 10852 | 17289 | 3407 | 0 | 8125 | 8124 | 8114 | 8133 | 8140 | 8148 |
| **697T** | **476** | 13743 | 15761 | 14017 | 14293 | 14294 | 14924 | 14909 | 13068 | 13919 | 13904 | 12413 | 13220 | 13231 | 13215 | 11751 | 11610 | 11613 | 10610 | 12054 | 17412 | 9901 | 8125 | 0 | 3 | 85 | 111 | 115 | 121 |
| **712T** | **476** | 13742 | 15760 | 14016 | 14292 | 14293 | 14923 | 14908 | 13067 | 13918 | 13903 | 12412 | 13219 | 13230 | 13214 | 11750 | 11609 | 11612 | 10609 | 12053 | 17411 | 9900 | 8124 | 3 | 0 | 84 | 110 | 114 | 120 |
| **18026** | **476** | 13726 | 15745 | 13998 | 14270 | 14271 | 14906 | 14891 | 13050 | 13899 | 13884 | 12398 | 13201 | 13212 | 13196 | 11731 | 11593 | 11596 | 10591 | 12042 | 17392 | 9892 | 8114 | 85 | 84 | 0 | 90 | 94 | 100 |
| **P.En250** | **476** | 13745 | 15769 | 14021 | 14293 | 14292 | 14931 | 14916 | 13073 | 13926 | 13911 | 12419 | 13224 | 13233 | 13219 | 11755 | 11612 | 11615 | 10622 | 12064 | 17416 | 9913 | 8133 | 111 | 110 | 90 | 0 | 116 | 122 |
| **11340** | **476** | 13754 | 15780 | 14028 | 14308 | 14309 | 14938 | 14923 | 13082 | 13933 | 13918 | 12424 | 13239 | 13250 | 13234 | 11761 | 11619 | 11624 | 10623 | 12065 | 17425 | 9918 | 8140 | 115 | 114 | 94 | 116 | 0 | 114 |
| **27149** | **858** | 13758 | 15782 | 14034 | 14309 | 14310 | 14950 | 14935 | 13086 | 13939 | 13924 | 12436 | 13237 | 13248 | 13232 | 11773 | 11627 | 11632 | 10631 | 12067 | 17424 | 9922 | 8148 | 121 | 120 | 100 | 122 | 114 | 0 |

**Table S2.** Acquired genes and chromosomal mutations conferring antibiotic resistance in the 28 analysed strains.

| Isolate[a] | ST | Country | Source | *n* | *optrA* variant[b] | Acquired resistance genes[c] | | | | | | | | | | | | | | | | | | | |
|---|---|---|---|---|---|---|---|---|---|---|---|---|---|---|---|---|---|---|---|---|---|---|---|---|---|
| | | | | | | Oxazolidinones | | | Phenicols | | Aminoglycosides | | | | | Macrolide/Lincosamide/Streptogramin B | | | | | Tetracyclines | | | | Trimethoprim |
| Enfs51 | 754 | MAL | PF | 12 | *optrA_7* | *optrA* | | | | *cat* | *aph(3')-III* | *aac(6')-aph(2'')* | | | *spc* | *lsa* (A) | *lsa* (E) | 2 Δ*erm* (B) | *erm* (A) | *lnu* (B) | *tet* (M) | *tet* (L) | | | |
| Enfs94 | 227 | MAL | HH | 1 | - | | | | | | | | | | | *lsa* (A) | | | | | | | | | |
| 728T | 859 | TUN | RC | 7 | *optrA_8* | *optrA* | | | *fexA* | | | | | | | *lsa* (A) | | | *erm* (A) | | *tet* (M) | *tet* (L) | *tet* (O) | | |
| 745E | 489 | COL | RC | 8 | *optrA_2* | *optrA* | | | *fexA* | | *aph(3')-III* | *aac(6')-aph(2'')* | | | | *lsa* (A) | | | | *lnu* (B) | Δ*tet* (M) | *tet* (L) | | | |
| N60443F | 489 | USA | RCA | 5 | *optrA_25* | *optrA* | | | *fexA* | | | | | | | *lsa* (A) | | 3 *erm* (B) | | | *tet* (M) | | | | |
| Iso4Bar | 474 | ESP | HP | 15 | *optrA_22* | Δ*optrA* | | | *fexA* | *cat* | *aph(3')-III* | *aac(6')-aph(2'')* | *ant(6)-Ia1* | *str* | *spc* | *lsa* (A) | | *erm* (B) | *erm* (A) | *lnu* (B) | *tet* (M) | *tet* (L) | | | Δ*dfrG* |
| Iso5Bar | 474 | ESP | HP | 13 | *optrA_23* | *optrA* | | | *fexA* | Δ*cat* | *aph(3')-III* | | *ant(6)-Ia1* | | *spc* | *lsa* (A) | | *erm* (B) | *erm* (A) | *lnu* (B) | *tet* (M) | *tet* (L) | | | Δ*dfrG* |
| Enfs85 | 755 | MAL | PF | 11 | *optrA_7* | *optrA* | | | *fexA* | | *aph(3')-III* | | Δ*ant(6)-Ia1* | *str* | *spc* | *lsa* (A) | | *erm* (B) | *erm* (A) | | *tet* (M) | *tet* (L) | | | |
| 12E | 59 | COL | RC | 7 | *optrA_2* | *optrA* | | | *fexA* | | | | | | | *lsa* (A) | | Δ*erm* (B) | | | *tet* (M) | *tet* (L) | | *tet* (K) | |
| 34E | 59 | COL | RC | 6 | *optrA_20* | *optrA* | | | *fexA* | | | | | | | *lsa* (A) | | *erm* (B) | | | *tet* (M) | *tet* (L) | | | |
| 12T | 86 | TUN | WW | 10 | ***optrA_2*** | *optrA* | | | *fexA* | | | *aac(6')-aph(2'')* | Δ*ant(6)-Ia1* | | *spc* | *lsa* (A) | | *erm* (B) | *erm* (A) | | *tet* (M) | | | | *dfrG* |
| Iso1Bar | 585 | ESP | HP | 8 | *optrA_3* | *optrA* | | | *fexA* | *cat* | | | | *str* | | *lsa* (A) | | *erm* (B) | | | *tet* (M) | *tet* (L) | | | |
| Iso2Bar | 585 | ESP | HP | 13 | *optrA_3* | *optrA* | | | *fexA* | *cat* | *aph(3')-III* | *aac(6')-aph(2'')* | *ant(6)-Ia* [#] | *str* | | *lsa* (A) | | *erm* (B) | | *lnu* (B) | *tet* (M) | *tet* (L) | | | Δ*dfrG* |
| Iso3Bar | 585 | ESP | HP | 8 | *optrA_3* | *optrA* | | | *fexA* | *cat* | | | | *str* | | *lsa* (A) | | *erm* (B) | | | *tet* (M) | *tet* (L) | | | |
| N48037F | 141 | USA | RP | 4 | ***optrA_26*** | *optrA* | | | *fexA* | | | | | | | *lsa* (A) | | | | | *tet* (M) | | | | |
| 612T | 21 | TUN | RC | 11 | ***optrA_7*** | *optrA* | | | *fexA* | *cat* | *aph(3')-III* | | | | *spc* | *lsa* (A) | | *erm* (B) | *erm* (A) | | *tet* (M) | *tet* (L) | | | *dfrG* |
| 13484 | 21 | CHI | HP | 10 | *optrA_5* | *optrA* | | | *fexA* | *cat* | | | | | *spc* | *lsa* (A) | | *erm* (B) | *erm* (A) | | *tet* (M) | *tet* (L) | | | *dfrG* |
| P.En218 | 765 | MAL | HH | 15 | *optrA_7* | *optrA* | | | *fexA* | *cat* | *aph(3')-III* | *aac(6')-aph(2'')* | *ant(6)-Ia* [#] | *str* | *spc* | *lsa* (A) | | *erm* (B) | *erm* (A) | *lnu* (B) | *tet* (M) | *tet* (L) | | | Δ*dfrG* |
| 16-372 | 480 | FRA | HP | 10 | *optrA_5* | *optrA* | | | *fexA* | | *aph(3')-III* | *aac(6')-aph(2'')* | | | *spc* | *lsa* (A) | *lsa* (E) | Δ*erm* (B) | *erm* (A) | *lnu* (B) | *tet* (M) | *tet* (L) | | | *dfrG* |
| P.En090 | 756 | MAL | PF | 13 | *optrA_7* | *optrA* | | | *fexA* | | *aph(3')-III* | *aac(6')-aph(2'')* | *ant(6)-Ia* [#] | | Δ*spc* | *lsa* (A) | | *erm* (B) | *erm* (A) | *lnu* (B) | *tet* (M) | *tet* (L) | | | *dfrG* |
| Efs 599 | 16 | USA | HH | 14 | *optrA_21* | *optrA* | *cfr1* | *poxtA* | | *cat* | *aph(3')-III* | *aac(6')-aph(2'')* | Δ*ant(6)-Ia1* | | Δ*spc* | *lsa* (A) | | *erm* (B) | *erm* (A) | | *tet* (M) | *tet* (L) | | | Δ*dfrG* |
| IBUN9046YE | 16 | COL | PF | 12 | *optrA_11* | *optrA* | | | *fexA* | Δ*cat* | *aph(3')-III* | *aac(6')-aph(2'')* | | | | *lsa* (A) | | Δ*erm* (B) | *erm* (A) | *lnu* (B) | *tet* (M) | *tet* (L) | | | *dfrG* |
| 697T | 476 | TUN | RC | 12 | ***optrA_5*** | *optrA* | | | *fexA* | *cat* | *aph(3')-III* | *aac(6')-aph(2'')* | | | *spc* | *lsa* (A) | | Δ*erm* (B) | *erm* (A) | | *tet* (M) | *tet* (L) | | | Δ*dfrG* |
| 712T | 476 | TUN | CF | 13 | ***optrA_5*** | *optrA* | | | *fexA* | *cat* | *aph(3')-III* | *aac(6')-aph(2'')* | *ant(6)-Ia1* | | *spc* | *lsa* (A) | | *erm* (B) | *erm* (A) | | *tet* (M) | *tet* (L) | | | Δ*dfrG* |
| 18026 | 476 | CHI | HP | 10 | *optrA_16* | *optrA* | | | *fexA* | *cat* | *aph(3')-III* | Δ*aac(6')-aph(2'')* | | | | *lsa* (A) | | *erm* (B) | *erm* (A) | | *tet* (M) | *tet* (L) | | | |
| P.En250 | 476 | MAL | PF | 14 | *optrA_24* | *optrA* | | | *fexA* | *cat* | *aph(3')-III* | *aac(6')-aph(2'')* | Δ*ant(6)-Ia1* | *str* | *spc* | *lsa* (A) | | *erm* (B) | *erm* (A) | | *tet* (M) | *tet* (L) | | | Δ*dfrG* |
| 11340 | 476 | CHI | HP | 14 | *optrA_8* | *optrA* | | | *fexA* | *cat* | *aph(3')-III* | *aac(6')-aph(2'')* | Δ*ant(6)-Ia1* | *str* | *spc* | *lsa* (A) | | *erm* (B) | *erm* (A) | | *tet* (M) | *tet* (L) | | | *dfrG* |
| 27149 | 858 | CHI | HP | 12 | *optrA_5* | *optrA* | | | *fexA* | *cat* | *aph(3')-III* | *aac(6')-aph(2'')* | | *str* | *spc* | *lsa* (A) | | Δ*erm* (B) | *erm* (A) | *lnu* (B) | | | | | *dfrG* |

Abbreviations: ST, sequence type; *n* , number of antibiotic resistance genes; CHI, China; COL, Colombia; ESP, Spain; FRA, France; MAL, Malaysia; TUN, Tunisia; USA, United States of America; CF, chicken feces; HH, healthy human; HP, hospitalized patient; PF, pig feces; RC,

[a]Isolates are ordered and separated according with their clustering in the phylogenetic tree (Figure 1).

[b]The *optrA* variants are given in comparison with the reference sequence of pE349 (KP399637.1). See supplementary Table S6 for classification details. *optrA* genes marked in bold are known to transfer, that in red to not transfer and in the remaining transfer is unknown.

[c]Acquired resistance genes are presented in different grey tones according with the highest homology they present with strain references retrieved in ResFinder (https://cge.cbs.dtu.dk/services/ResFinder/). [#]These isolates additionally contained *Δant(6)-Ia1* .

Homology: 100% ≥99%<100% ≥97%<99% ≥95%<97%

[d]The strain reference for linezolid resistance mutations was *E. faecalis* V583. Underlined mutations are also described by Bender *et al* (9).

[e]23S rRNA synonymous mutations are assigned (syn) while non-synonymous 23S rRNA mutations were not previously described (G388A/D130N, G377A/R126K, G693A/M231I, G1457A/S486N, G2548A/G850S, G116A/R39Q, C307T/P102S).

**Table S3.** Extended virulence gene profiles present in the 28 analysed strains.

| Isolate[a] | ST | Country | Source | *n* | Secreted factors for tissue damage | | | | | | | | | Adhesins and Biofilm-associated genes | | | | | | | Survival | | General Stress | | Membrane Metalloprotease |
|---|---|---|---|---|---|---|---|---|---|---|---|---|---|---|---|---|---|---|---|---|---|---|---|---|---|
| Enfs51 | 754 | MAL | PF | 38 | | | | | *gelE* | *fsrB* | *hylA* | | | | *efaAfs* | *ebpA* | *ebpB* | | *srtA* | *efbA* | *elrA* | *tpx* | *gls24* | *glsB* | *eep* |
| Enfs94 | 227 | MAL | HH | 37 | | | | | *gelE* | *fsrB* | Δ*hylA* | | | | *efaAfs* | *ebpA* | *ebpB* | | *srtA* | *efbA* | *elrA* | *tpx* | *gls24* | *glsB* | *eep* |
| 728T | 859 | TUN | RC | 52 | *cylA* | | *cylL* | *cylM* | *gelE* | *fsrB* | | *hylB* | *agg* | *ace* | *efaAfs* | *ebpA* | | *ebpC* | *srtA* | *efbA* | *elrA* | *tpx* | *gls24* | *glsB* | *eep* |
| 745E | 489 | COL | RC | 38 | | | | | Δ*gelE* | *fsrB* | ↑*hylA* | *hylB* | | *ace* | *efaAfs* | *ebpA* | | *ebpC* | *srtA* | *efbA* | *elrA* | *tpx* | *gls24* | *glsB* | *eep* |
| N60443F | 489 | USA | RCA | 41 | | | | | *gelE* | *fsrB* | *hylA* | *hylB* | | *ace* | *efaAfs* | *ebpA* | *ebpB* | *ebpC* | *srtA* | *efbA* | *elrA* | *tpx* | *gls24* | *glsB* | *eep* |
| Iso4Bar | 474 | ESP | HP | 42 | | | | | Δ*gelE* | *fsrB* | ↑*hylA* | *hylB* | Δ*agg* | *ace* | *efaAfs* | *ebpA* | *ebpB* | *ebpC* | *srtA* | *efbA* | *elrA* | *tpx* | *gls24* | *glsB* | *eep* |
| Iso5Bar | 474 | ESP | HP | 42 | | | | | Δ*gelE* | *fsrB* | ↑*hylA* | *hylB* | Δ*agg* | *ace* | *efaAfs* | *ebpA* | *ebpB* | *ebpC* | *srtA* | *efbA* | *elrA* | *tpx* | *gls24* | *glsB* | *eep* |
| Enfs85 | 755 | MAL | PF | 38 | | | | | | | | *hylB* | Δ*agg* | | *efaAfs* | *ebpA* | *ebpB* | *ebpC* | *srtA* | *efbA* | *elrA* | *tpx* | *gls24* | *glsB* | *eep* |
| 12E | 59 | COL | RC | 40 | | | | | Δ*gelE* | *fsrB* | Δ*hylA* | *hylB* | Δ*agg* | *ace* | *efaAfs* | *ebpA* | | *ebpC* | *srtA* | *efbA* | *elrA* | *tpx* | *gls24* | *glsB* | *eep* |
| 34E | 59 | COL | RC | 39 | | | | | Δ*gelE* | *fsrB* | Δ*hylA* | *hylB* | | *ace* | *efaAfs* | *ebpA* | | *ebpC* | *srtA* | *efbA* | *elrA* | *tpx* | *gls24* | *glsB* | *eep* |
| 12T | 86 | TUN | WW | 47 | | | | | *gelE* | *fsrB* | ↑*hylA* | *hylB* | *agg* | *ace* | | *ebpA* | | *ebpC* | *srtA* | *efbA* | *elrA* | *tpx* | *gls24* | *glsB* | *eep* |
| Iso1Bar | 585 | ESP | HP | 50 | *cylA* | | *cylL* | *cylM* | *gelE* | *fsrB* | | | | *ace* | *efaAfs* | *ebpA* | | *ebpC* | *srtA* | *efbA* | *elrA* | *tpx* | *gls24* | *glsB* | *eep* |
| Iso2Bar | 585 | ESP | HP | 50 | *cylA* | | *cylL* | *cylM* | *gelE* | *fsrB* | | | | *ace* | *efaAfs* | *ebpA* | | *ebpC* | *srtA* | *efbA* | *elrA* | *tpx* | *gls24* | *glsB* | *eep* |
| Iso3Bar | 585 | ESP | HP | 50 | *cylA* | | *cylL* | *cylM* | *gelE* | *fsrB* | | | | *ace* | *efaAfs* | *ebpA* | | *ebpC* | *srtA* | *efbA* | *elrA* | *tpx* | *gls24* | *glsB* | *eep* |
| N48037F | 141 | USA | RP | 54 | *cylA* | *cylB* | *cylL* | *cylM* | *gelE* | *fsrB* | *hylA* | *hylB* | *agg* | *ace* | *efaAfs* | *ebpA* | *ebpB* | *ebpC* | *srtA* | *efbA* | *elrA* | *tpx* | *gls24* | *glsB* | *eep* |
| 612T | 21 | TUN | RC | 40 | | | | | *gelE* | *fsrB* | ↑*hylA* | *hylB* | | *ace* | *efaAfs* | *ebpA* | | *ebpC* | *srtA* | *efbA* | *elrA* | *tpx* | *gls24* | *glsB* | *eep* |
| 13484 | 21 | CHI | HP | 41 | | | | | *gelE* | *fsrB* | *hylA* | *hylB* | | *ace* | *efaAfs* | *ebpA* | *ebpB* | *ebpC* | *srtA* | *efbA* | *elrA* | *tpx* | *gls24* | *glsB* | *eep* |
| P.En218 | 765 | MAL | HH | 39 | | | | | *gelE* | *fsrB* | Δ*hylA* | | | *ace* | *efaAfs* | *ebpA* | *ebpB* | | *srtA* | *efbA* | *elrA* | *tpx* | *gls24* | *glsB* | *eep* |
| 16-372 | 480 | FRA | HP | 38 | | | | | | | ↑*hylA* | | *agg* | *ace* | *efaAfs* | *ebpA* | *ebpB* | *ebpC* | *srtA* | *efbA* | *elrA* | *tpx* | *gls24* | *glsB* | *eep* |
| P.En090 | 756 | MAL | PF | 42 | | | | | *gelE* | *fsrB* | Δ*hylA* | *hylB* | *agg* | *ace* | *efaAfs* | *ebpA* | *ebpB* | | *srtA* | *efbA* | *elrA* | *tpx* | *gls24* | *glsB* | *eep* |
| Efs 599 | 16 | USA | HH | 50 | *cylA* | | *cylL* | *cylM* | *gelE* | *fsrB* | ↑*hylA* | | | *ace* | *efaAfs* | *ebpA* | | *ebpC* | *srtA* | *efbA* | *elrA* | *tpx* | *gls24* | *glsB* | *eep* |
| IBUN9046YE | 16 | COL | PF | 52 | *cylA* | | *cylL* | *cylM* | *gelE* | *fsrB* | ↑*hylA* | | *agg* | *ace* | *efaAfs* | *ebpA* | | *ebpC* | *srtA* | *efbA* | *elrA* | *tpx* | *gls24* | *glsB* | *eep* |
| 697T | 476 | TUN | RC | 47 | | | | | | | ↑*hylA* | | Δ*agg* | *ace* | *efaAfs* | *ebpA* | *ebpB* | *ebpC* | *srtA* | *efbA* | *elrA* | *tpx* | *gls24* | *glsB* | *eep* |
| 712T | 476 | TUN | CF | 46 | | | | | | | ↑*hylA* | | Δ*agg* | *ace* | *efaAfs* | *ebpA* | | *ebpC* | *srtA* | *efbA* | *elrA* | *tpx* | *gls24* | *glsB* | *eep* |
| 18026 | 476 | CHI | HP | 46 | | | | | | | ↑*hylA* | | | *ace* | *efaAfs* | *ebpA* | *ebpB* | *ebpC* | *srtA* | *efbA* | *elrA* | *tpx* | *gls24* | *glsB* | *eep* |
| P.En250 | 476 | MAL | PF | 46 | | | | | | | ↑*hylA* | | | *ace* | *efaAfs* | *ebpA* | | *ebpC* | *srtA* | *efbA* | *elrA* | *tpx* | *gls24* | *glsB* | *eep* |
| 11340 | 476 | CHI | HP | 47 | | | | | | | ↑*hylA* | | Δ*agg* | *ace* | *efaAfs* | *ebpA* | *ebpB* | *ebpC* | *srtA* | *efbA* | *elrA* | *tpx* | *gls24* | *glsB* | *eep* |
| 27149 | 858 | CHI | HP | 47 | | | | | | | ↑*hylA* | | Δ*agg* | *ace* | *efaAfs* | *ebpA* | *ebpB* | *ebpC* | *srtA* | *efbA* | *elrA* | *tpx* | *gls24* | *glsB* | *eep* |

| Isolate[a] | Capsule and Cell-wall Components | | | | | | | | | | | | | | | | |
|---|---|---|---|---|---|---|---|---|---|---|---|---|---|---|---|---|---|
| Enfs51 | | | | | | | | | | *epaA* | *epaB* | *epaC* | *epaD* | *epaE* | *epaF* | *epaG* | *epaH* |
| Enfs94 | | | | | | | | | | *epaA* | *epaB* | *epaC* | *epaD* | *epaE* | *epaF* | *epaG* | *epaH* |
| 728T | *cpsC* | *cpsD* | *cpsE* | ↑*cpsF* | ↑*cpsG* | *cpsH* | *cpsI* | *cpsJ* | *cpsK* | *epaA* | *epaB* | *epaC* | *epaD* | *epaE* | *epaF* | *epaG* | *epaH* |
| 745E | | | | | | | | | | *epaA* | *epaB* | *epaC* | *epaD* | *epaE* | *epaF* | *epaG* | *epaH* |
| N60443F | | | | | | | | | | *epaA* | *epaB* | *epaC* | *epaD* | *epaE* | *epaF* | *epaG* | *epaH* |
| Iso4Bar | | | | | | | | | | *epaA* | *epaB* | *epaC* | *epaD* | *epaE* | *epaF* | *epaG* | *epaH* |
| Iso5Bar | | | | | | | | | | *epaA* | *epaB* | *epaC* | *epaD* | *epaE* | *epaF* | *epaG* | *epaH* |
| Enfs85 | | | | | | | | | | *epaA* | *epaB* | *epaC* | *epaD* | *epaE* | *epaF* | *epaG* | *epaH* |
| 12E | | | | | | | | | | *epaA* | *epaB* | *epaC* | *epaD* | *epaE* | *epaF* | *epaG* | *epaH* |
| 34E | | | | | | | | | | *epaA* | *epaB* | *epaC* | *epaD* | *epaE* | *epaF* | *epaG* | *epaH* |
| 12T | *cpsC* | *cpsD* | *cpsE* | | ↑*cpsG* | *cpsH* | *cpsI* | *cpsJ* | *cpsK* | *epaA* | *epaB* | *epaC* | *epaD* | *epaE* | *epaF* | *epaG* | *epaH* |
| Iso1Bar | *cpsC* | *cpsD* | *cpsE* | ↑*cpsF* | ↑*cpsG* | *cpsH* | *cpsI* | *cpsJ* | *cpsK* | *epaA* | *epaB* | *epaC* | *epaD* | *epaE* | *epaF* | *epaG* | *epaH* |
| Iso2Bar | *cpsC* | *cpsD* | *cpsE* | ↑*cpsF* | ↑*cpsG* | *cpsH* | *cpsI* | *cpsJ* | *cpsK* | *epaA* | *epaB* | *epaC* | *epaD* | *epaE* | *epaF* | *epaG* | *epaH* |
| Iso3Bar | *cpsC* | *cpsD* | *cpsE* | ↑*cpsF* | ↑*cpsG* | *cpsH* | *cpsI* | *cpsJ* | *cpsK* | *epaA* | *epaB* | *epaC* | *epaD* | *epaE* | *epaF* | *epaG* | *epaH* |
| N48037F | *cpsC* | *cpsD* | ↑*cpsE* | | ↑*cpsG* | *cpsH* | *cpsI* | *cpsJ* | *cpsK* | *epaA* | *epaB* | *epaC* | *epaD* | *epaE* | *epaF* | *epaG* | *epaH* |
| 612T | | | | | | | | | | *epaA* | *epaB* | *epaC* | *epaD* | *epaE* | *epaF* | *epaG* | *epaH* |
| 13484 | | | | | | | | | | *epaA* | *epaB* | *epaC* | *epaC* | *epaE* | *epaF* | *epaG* | *epaH* |
| P.En218 | | | | | | | | | | *epaA* | *epaB* | *epaC* | *epaD* | *epaE* | *epaF* | *epaG* | *epaH* |
| 16-372 | | | | | | | | | | *epaA* | *epaB* | *epaC* | *epaD* | *epaE* | *epaF* | *epaG* | *epaH* |
| P.En090 | | | | | | | | | | *epaA* | *epaB* | *epaC* | *epaD* | *epaE* | *epaF* | *epaG* | *epaH* |
| Efs 599 | *cpsC* | *cpsD* | *cpsE* | ↑*cpsF* | ↑*cpsG* | *cpsH* | *cpsI* | *cpsJ* | *cpsK* | *epaA* | *epaB* | *epaC* | *epaD* | *epaE* | *epaF* | *epaG* | *epaH* |
| IBUN9046YE | *cpsC* | *cpsD* | *cpsE* | ↑*cpsF* | ↑*cpsG* | *cpsH* | *cpsI* | *cpsJ* | *cpsK* | *epaA* | *epaB* | *epaC* | *epaD* | *epaE* | *epaF* | *epaG* | *epaH* |
| 697T | *cpsC* | *cpsD* | *cpsE* | ↑*cpsF* | ↑*cpsG* | *cpsH* | *cpsI* | *cpsJ* | *cpsK* | *epaA* | *epaB* | *epaC* | *epaD* | *epaE* | *epaF* | *epaG* | *epaH* |
| 712T | *cpsC* | *cpsD* | *cpsE* | ↑*cpsF* | ↑*cpsG* | *cpsH* | *cpsI* | *cpsJ* | *cpsK* | *epaA* | *epaB* | *epaC* | *epaD* | *epaE* | *epaF* | *epaG* | *epaH* |
| 18026 | *cpsC* | *cpsD* | *cpsE* | ↑*cpsF* | ↑*cpsG* | *cpsH* | *cpsI* | *cpsJ* | *cpsK* | *epaA* | *epaB* | *epaC* | *epaD* | *epaE* | *epaF* | *epaG* | *epaH* |
| P.En250 | *cpsC* | *cpsD* | *cpsE* | ↑*cpsF* | ↑*cpsG* | *cpsH* | *cpsI* | *cpsJ* | *cpsK* | *epaA* | *epaB* | *epaC* | *epaD* | *epaE* | *epaF* | *epaG* | *epaH* |
| 11340 | *cpsC* | *cpsD* | *cpsE* | ↑*cpsF* | ↑*cpsG* | *cpsH* | *cpsI* | *cpsJ* | *cpsK* | *epaA* | *epaB* | *epaC* | *epaD* | *epaE* | *epaF* | *epaG* | *epaH* |
| 27149 | *cpsC* | *cpsD* | *cpsE* | ↑*cpsF* | ↑*cpsG* | *cpsH* | *cpsI* | *cpsJ* | *cpsK* | *epaA* | *epaB* | *epaC* | *epaD* | *epaE* | *epaF* | *epaG* | *epaH* |

| Isolate[a] | Capsule and Cell-wall Components (cont.) | | | | | | | | | | | | Sex pheromones | | | | PAI | |
|---|---|---|---|---|---|---|---|---|---|---|---|---|---|---|---|---|---|---|
| Enfs51 | Δ*epaI* | Δ*epaJ* | *epaK* | *epaL* | *epaM* | *epaN* | *epaO* | *epaP* | *epaQ* | *epaR* | *bgsA* | *bgsB* | *cad* | *camE* | *cCF10* | *cOB1* | *cbh* | |
| Enfs94 | Δ*epaI* | Δ*epaJ* | *epaK* | *epaL* | *epaM* | *epaN* | *epaO* | *epaP* | *epaQ* | *epaR* | *bgsA* | *bgsB* | *cad* | *camE* | *cCF10* | *cOB1* | | |
| 728T | Δ*epaI* | Δ*epaJ* | *epaK* | *epaL* | *epaM* | *epaN* | *epaO* | *epaP* | *epaQ* | *epaR* | *bgsA* | *bgsB* | *cad* | *camE* | *cCF10* | *cOB1* | | *ef0571* |
| 745E | *epaI* | Δ*epaJ* | *epaK* | *epaL* | | *epaN* | *epaO* | *epaP* | *epaQ* | *epaR* | *bgsA* | *bgsB* | *cad* | *camE* | *cCF10* | *cOB1* | | |
| N60443F | *epaI* | Δ*epaJ* | *epaK* | *epaL* | *epaM* | *epaN* | *epaO* | *epaP* | *epaQ* | *epaR* | *bgsA* | *bgsB* | *cad* | *camE* | *cCF10* | *cOB1* | *cbh* | |
| Iso4Bar | Δ*epaI* | Δ*epaJ* | *epaK* | *epaL* | *epaM* | *epaN* | *epaO* | *epaP* | *epaQ* | *epaR* | *bgsA* | *bgsB* | *cad* | *camE* | *cCF10* | *cOB1* | *cbh* | |
| Iso5Bar | Δ*epaI* | Δ*epaJ* | *epaK* | *epaL* | *epaM* | *epaN* | *epaO* | *epaP* | *epaQ* | *epaR* | *bgsA* | *bgsB* | *cad* | *camE* | *cCF10* | *cOB1* | *cbh* | |
| Enfs85 | Δ*epaI* | Δ*epaJ* | *epaK* | *epaL* | *epaM* | *epaN* | *epaO* | *epaP* | *epaQ* | *epaR* | *bgsA* | *bgsB* | *cad* | *camE* | *cCF10* | *cOB1* | *cbh* | |
| 12E | Δ*epaI* | Δ*epaJ* | *epaK* | *epaL* | | *epaN* | *epaO* | *epaP* | *epaQ* | *epaR* | *bgsA* | *bgsB* | *cad* | *camE* | *cCF10* | *cOB1* | *cbh* | |
| 34E | Δ*epaI* | Δ*epaJ* | *epaK* | *epaL* | | *epaN* | *epaO* | *epaP* | *epaQ* | *epaR* | *bgsA* | *bgsB* | *cad* | *camE* | *cCF10* | *cOB1* | *cbh* | |
| 12T | Δ*epaI* | Δ*epaJ* | *epaK* | *epaL* | *epaM* | *epaN* | *epaO* | *epaP* | *epaQ* | *epaR* | *bgsA* | *bgsB* | *cad* | *camE* | *cCF10* | *cOB1* | | |
| Iso1Bar | Δ*epaI* | Δ*epaJ* | *epaK* | *epaL* | *epaM* | *epaN* | *epaO* | *epaP* | *epaQ* | *epaR* | *bgsA* | *bgsB* | *cad* | *camE* | *cCF10* | *cOB1* | *cbh* | |
| Iso2Bar | Δ*epaI* | Δ*epaJ* | *epaK* | *epaL* | *epaM* | *epaN* | *epaO* | *epaP* | *epaQ* | *epaR* | *bgsA* | *bgsB* | *cad* | *camE* | *cCF10* | *cOB1* | *cbh* | |
| Iso3Bar | Δ*epaI* | Δ*epaJ* | *epaK* | *epaL* | *epaM* | *epaN* | *epaO* | *epaP* | *epaQ* | *epaR* | *bgsA* | *bgsB* | *cad* | *camE* | *cCF10* | *cOB1* | *cbh* | |
| N48037F | Δ*epaI* | *epaJ* | *epaK* | *epaL* | *epaM* | *epaN* | *epaO* | *epaP* | *epaQ* | *epaR* | *bgsA* | *bgsB* | *cad* | *camE* | *cCF10* | *cOB1* | *cbh* | |
| 612T | Δ*epaI* | Δ*epaJ* | *epaK* | *epaL* | *epaM* | *epaN* | *epaO* | *epaP* | *epaQ* | *epaR* | *bgsA* | *bgsB* | *cad* | *camE* | *cCF10* | *cOB1* | *cbh* | |
| 13484 | Δ*epaI* | Δ*epaJ* | *epaK* | *epaL* | *epaM* | *epaN* | *epaO* | *epaP* | *epaQ* | *epaR* | *bgsA* | *bgsB* | *cad* | *camE* | *cCF10* | *cOB1* | *cbh* | |
| P.En218 | Δ*epaI* | Δ*epaJ* | *epaK* | *epaL* | *epaM* | *epaN* | *epaO* | *epaP* | *epaQ* | *epaR* | *bgsA* | *bgsB* | *cad* | *camE* | *cCF10* | *cOB1* | *cbh* | |
| 16-372 | Δ*epaI* | Δ*epaJ* | *epaK* | *epaL* | *epaM* | *epaN* | *epaO* | *epaP* | *epaQ* | *epaR* | *bgsA* | *bgsB* | *cad* | *camE* | *cCF10* | *cOB1* | | |
| P.En090 | Δ*epaI* | Δ*epaJ* | *epaK* | *epaL* | *epaM* | *epaN* | *epaO* | *epaP* | *epaQ* | *epaR* | *bgsA* | *bgsB* | *cad* | *camE* | *cCF10* | *cOB1* | *cbh* | *ef0571* |
| Efs 599 | Δ*epaI* | Δ*epaJ* | *epaK* | *epaL* | *epaM* | *epaN* | *epaO* | *epaP* | *epaQ* | *epaR* | *bgsA* | *bgsB* | *cad* | *camE* | *cCF10* | *cOB1* | | |
| IBUN9046YE | Δ*epaI* | Δ*epaJ* | *epaK* | *epaL* | *epaM* | *epaN* | *epaO* | *epaP* | *epaQ* | *epaR* | *bgsA* | *bgsB* | *cad* | *camE* | *cCF10* | *cOB1* | *cbh* | |
| 697T | Δ*epaI* | Δ*epaJ* | *epaK* | *epaL* | *epaM* | *epaN* | *epaO* | *epaP* | *epaQ* | *epaR* | *bgsA* | *bgsB* | *cad* | *camE* | *cCF10* | *cOB1* | | |
| 712T | Δ*epaI* | Δ*epaJ* | *epaK* | *epaL* | *epaM* | *epaN* | *epaO* | *epaP* | *epaQ* | *epaR* | *bgsA* | *bgsB* | *cad* | *camE* | *cCF10* | *cOB1* | | |
| 18026 | Δ*epaI* | Δ*epaJ* | *epaK* | *epaL* | *epaM* | *epaN* | *epaO* | *epaP* | *epaQ* | *epaR* | *bgsA* | *bgsB* | *cad* | *camE* | *cCF10* | *cOB1* | | |
| P.En250 | Δ*epaI* | Δ*epaJ* | *epaK* | *epaL* | *epaM* | *epaN* | *epaO* | *epaP* | *epaQ* | *epaR* | *bgsA* | *bgsB* | *cad* | *camE* | *cCF10* | *cOB1* | *cbh* | |
| 11340 | Δ*epaI* | Δ*epaJ* | *epaK* | *epaL* | *epaM* | *epaN* | *epaO* | *epaP* | *epaQ* | *epaR* | *bgsA* | *bgsB* | *cad* | *camE* | *cCF10* | *cOB1* | | |
| 27149 | Δ*epaI* | Δ*epaJ* | *epaK* | *epaL* | *epaM* | *epaN* | *epaO* | *epaP* | *epaQ* | *epaR* | *bgsA* | *bgsB* | *cad* | *camE* | *cCF10* | *cOB1* | | |

Abbreviations: ST, sequence type; *n*, number of virulence genes; CHI, China; COL, Colombia; ESP, Spain; FRA, France; MAL, Malaysia; TUN, Tunisia; USA, United States of America; CF, chicken feces; HH, healthy human; HP, hospitalized patient; PF, pig feces; RC, retail chicken; RCA, retail cattle; RP, retail pig; WW, wastewat

[a]Isolates are ordered and separated according with their clustering in the phylogenetic tree (Figure 1).

The virulence genes are presented in different grey tones according with the highest homology they present with strain references retrieved in ResFinder (https://cge.cbs.dtu.dk/services/ResFinder/) or *E. faecalis* V583 or OG1RF strains in the case of the virulence genes obtained from our in-house database. Truncated forms are represented with Δ and ↑ indicates genes with higher lengths.

Homology: 100%; ≥99%<100%; ≥97%<99%; ≥95%<97%; ≥90%<95%

**Table S4.** Prophages present in the 28 analysed strains.

| Isolate[a] | ST | Country | Source | *n* | **Prophages[b]** | | | | | | | | | | | | | | | | | | | |
|---|---|---|---|---|---|---|---|---|---|---|---|---|---|---|---|---|---|---|---|---|---|---|---|---|
| | | | | | ***Enterococcus*** | | | | | | ***Staphylococcus*** | ***Streptococcus*** | | ***Listeria*** | | | ***Lactobacillus*** | ***Paenibacillus*** | ***Bacillus*** | | | ***Lactococcus*** | ***Enterobacteria*** | ***Prochlorococcus*** |
| | | | | | **phiFL1A** | **vB_IME197** | **phiEf11** | **phiFL2A** | **phiFL3A** | **phiEF24C** | **SPbeta-like** | **EJ-1** | **T12** | **LP-101** | **LP-B025** | **LP-B054** | **PLE2** | **Diva-like** | **Shanette** | **phage-G** | **SPbeta** | **phiL47** | **phi92** | **P-SSM2** |
| Enfs51 | 754 | MAL | PF | 1 | | | | | | | | | | | | | | | | phage-G | | | | |
| Enfs94 | 227 | MAL | HH | 0 | | | | | | | | | | | | | | | | | | | | |
| 728T | 859 | TUN | RC | 3 | phiFL1A | vB_IME197 | | | | | | | | | | | | Diva-like | | | | | | |
| 745E | 489 | COL | RC | 1 | | | | | | | | | | | | | | | Shanette | | | | | |
| N60443F | 489 | USA | RCA | 2 | | vB_IME197 | | | | | | | | | | | | Diva-like | | | | | | |
| Iso4Bar | 474 | ESP | HP | 1 | | vB_IME197 | | | | | | | | | | | | | | | | | | |
| Iso5Bar | 474 | ESP | HP | 1 | | | | | | | | | | | | | | Diva-like | | | | | | |
| Enfs85 | 755 | MAL | PF | 1 | | | | | | | | | T12 | | | | | | | | | phiL47 | | |
| 12E | 59 | COL | RC | 1 | | | | | | | | | | | | | | | | | | | | P-SSM2 |
| 34E | 59 | COL | RC | 0 | | | | | | | | | | | | | | | | | | | | |
| 12T | 86 | TUN | WW | 2 | | **vB_IME197** | | | | | | | | **LP-101** | | | | | | | | | | |
| Iso1Bar | 585 | ESP | HP | 1 | **phiFL1A** | | | | | | | | | | | | | | | | | | | |
| Iso2Bar | 585 | ESP | HP | 1 | **phiFL1A** | | | | | | | | | | | | | | | | | | | |
| Iso3Bar | 585 | ESP | HP | 1 | **phiFL1A** | | | | | | | | | | | | | | | | | | | |
| N48037F | 141 | USA | RP | 4 | phiFL1A | vB_IME197 | | phiFL2A | **phiFL3A** | | | | | | | | | | | | | | | |
| 612T | 21 | TUN | RC | 1 | | **vB_IME197** | | | | | | | | | | | | | | | | | | |
| 13484 | 21 | CHI | HP | 3 | | vB_IME197 | | | | phiEF24C | | | | **LP-101** | | | | | | | | | | |
| P.En218 | 765 | MAL | HH | 1 | **phiFL1A** | | | | | | | | | | | | | | | | | | | |
| 16-372 | 480 | FRA | HP | 4 | **phiFL1A** | vB_IME197 | | | | | SPbeta-like | | | | **LP-B025** | | | | | | | | | |
| P.En090 | 756 | MAL | PF | 3 | **phiFL1A** | | | | | | | | | | | | | Diva-like | | | | phiL47 | | |
| Efs 599 | 16 | USA | HH | 2 | | | | | | | | EJ-1 | | | | | | | | | | | phi92 | |
| IBUN9046YE | 16 | COL | PF | 1 | **phiFL1A** | | | | | | | | | | | | | | | | | | | |
| 697T | 476 | TUN | RC | 7 | phiFL1A | **vB_IME197** | **phiEf11** | | phiFL3A | | | | | | | LP-B054 | PLE2 | | | | SPbeta | | | |
| 712T | 476 | TUN | CF | 1 | | | **phiEf11** | | | | | | | | | | | | | | | | | |
| 18026 | 476 | CHI | HP | 3 | **phiFL1A** | vB_IME197 | | | | | | | | | | | | | | | | | phi92 | |
| P.En250 | 476 | MAL | PF | 2 | | | | | | | | EJ-1 | | | | | | Diva-like | | | | | | |
| 11340 | 476 | CHI | HP | 3 | | vB_IME197 | | | | | | | | **LP-101** | | LP-B054 | | | | | | | | |
| 27149 | 858 | CHI | HP | 3 | **phiFL1A** | vB_IME197 | | | | | | | | | | | | | | | | | phi92 | |

Abbreviations: ST, sequence type; *n*, number of prophages; CHI, China; COL, Colombia; ESP, Spain; FRA, France; MAL, Malaysia; TUN, Tunisia; USA, United States of America; CF, chicken feces; HH, healthy human; HP, hospitalized patient; PF, pig feces; RC, retail chicken; RCA, retail cattle; RP, retail pig; WW, wastewaters.

[a]Isolates are ordered and separated according with their clsutering in the phylogenetic tree (Figure 1).

[b]Prophages sequences were identified by using the PHASTER web tool and those represented in bold are intact: phiFL1A (*E. faecalis* phage phiFL1A, NC_013646), vB_IME197 (*E. faecalis* phage vB_EfaS_IME197, NC_028671), phiEf11 (*E. faecalis* phage phiEf11, NC_013696), phiFL2A (*E. faecalis* phage phiFL2A, NC_013643), phiFL3A (*E. faecalis* phage phiFL3A, NC_013648), phiEF24C (*E. faecalis* phage phiEF24C, NC_009904), SPbeta-like (*Staphylococcus epidermidis* phage SPbeta-like, NC_029119), EJ-1 (*Streptococcus pneumoniae* phage EJ-1, NC_005294), T12 (*Streptococcus* phage T12, NC_028700), LP-101 (*Listeria* phage LP-101, NC_024387), LP-B025 (*Listeria* phage B025, NC_009812), LP-B054 (*Listeria* phage B054, NC_009813), PLE2 (*Lactobacillus casei* phage Lactob_PLE2, NC_031036), Diva-like (*Paenibacillus larvae* phage Diva, NC_028788), Shanette (*Bacillus cereus* phage Shanette, NC_028983), phage-G (*Bacillus* phage G, NC_023719), SPbeta (*Bacillus subtilis* phage SPBc2, NC_001884), phiL47 (*Lactococcus* phage phiL47, NC_023574), phi92 (*Enterobacteria* phage phi92, NC_023693), P-SSM2 (*Prochlorococcus* phage P-SSM2, NC_006883). Enfs94 and 34E strains did not carry prophage sequences.

**Table S5.** Replication initiation genes present in the 28 analysed strains.

| Isolate[a] | ST | Country | Source | *n* | Rolling Circle Replicating (RCR)[b] | | | | | Theta replicating[b] | | | | | | | | | | | | | | |
|---|---|---|---|---|---|---|---|---|---|---|---|---|---|---|---|---|---|---|---|---|---|---|---|---|
| | | | | | Rep trans | | | Rep1 | Rep2 | RepA_N | | | | | | Inc18 | | | | Rep3 | | | | |
| | | | | | *Enterococcus* | *Streptococcus* | *Staphylococcus* | *Staphylococcus* | *Lactococcus* | *Enterococcus* | | | | | | *Enterococcus* | | | *Streptococcus* | *Enterococcus* | | | | |
| Enfs51 | 754 | MAL | PF | 3 | rep$_{pDO1}$ | | | | | | | Δrep$_{pTEF1}$ | | rep$_{pTW9}$ | | | | | | | | | | |
| Enfs94 | 227 | MAL | HH | 0 | | | | | | | | | | | | | | | | | | | | |
| 728T | 859 | TUN | RC | 2 | rep$_{pDO1}$ | | | | | | | | | Δrep$_{pTW9}$ | | | | | | | | | | |
| 745E | 489 | COL | RC | 3 | rep$_{pDO1}$ | | | | | rep$_{EF62pC}$ | | | | | | | | rep$_{pTW9}$ | | | | | rep$_{pE394}$ | |
| N60443F | 489 | USA | RCA | 1 | | | | | | | | | | | | | | | | | | | rep$_{pE394}$ | |
| Iso4Bar | 474 | ESP | HP | 5 | rep$_{pDO1}$ | | | | | | | Δrep$_{pTEF1}$ | rep$_{pTEF2}$ | | | | | | | rep$_{pMBB1}$ | rep$_{pS86}$ | | | |
| Iso5Bar | 474 | ESP | HP | 3 | rep$_{pDO1}$ | | | | | | rep$_{pEJ97-1}$ | Δrep$_{pTEF1}$ | | | | | | | | | | | | |
| Enfs85 | 755 | MAL | PF | 5 | rep$_{pDO1}$ | rep$_{pGB354}$ | | | | | rep$_{pEJ97-1}$ | Δrep$_{pTEF1}$ | | | | | rep$_{pTEF3}$ | | | | | | | |
| 12E | 59 | COL | RC | 4 | rep$_{pDO1}$ | | rep$_{SAP093A}$ | | | rep$_{EF62pC}$ | | | | | | | rep$_{pTEF3}$ | | | | | | rep$_{pE394}$ | |
| 34E | 59 | COL | RC | 2 | rep$_{pDO1}$ | | | | | rep$_{EF62pC}$ | | | | | | | | | | | | | rep$_{pE394}$ | |
| 12T | 86 | TUN | WW | 3 | rep$_{pDO1}$ | | | | | | | | | | | rep$_{pRE25}$ | | | | | | rep$_{pE1_13}$ | | |
| Iso1Bar | 585 | ESP | HP | 3 | rep$_{pDO1}$ | rep$_{pGB354}$ | | | | | | Δrep$_{pTEF1}$ | | | | | | | | | | | | |
| Iso2Bar | 585 | ESP | HP | 3 | rep$_{pDO1}$ | rep$_{pGB354}$ | | | | | | Δrep$_{pTEF1}$ | | | | | | | | | | | | |
| Iso3Bar | 585 | ESP | HP | 3 | rep$_{pDO1}$ | rep$_{pGB354}$ | | | | | | Δrep$_{pTEF1}$ | | | | | | | | | | | | |
| N48037F | 141 | USA | RP | 4 | rep$_{pDO1}$ | | | | | | | | | rep$_{pTW9}$ | rep$_{pCF10}$ | | | | | | | | rep$_{pE394}$ | |
| 612T | 21 | TUN | RC | 2 | | | | rep$_{pKKS627}$ | | | | | | ↑rep$_{pTW9}$ | | | | | | | | | | |
| 13484 | 21 | CHI | HP | 4 | rep$_{pDO1}$ | | | rep$_{pKKS627}$ | | | | | | rep$_{pTW9}$ | | | | rep$_{pTW9}$ | | | | | | |
| P.En218 | 765 | MAL | HH | 4 | rep$_{pDO1}$ | rep$_{pGB354}$ | | | | | | Δrep$_{pTEF1}$ | | | | | rep$_{pTEF3}$ | | | | | | | |
| 16-372 | 480 | FRA | HP | 1 | rep$_{pDO1}$ | | | | | | | | | | | | | | | | | | | |
| P.En090 | 756 | MAL | PF | 3 | rep$_{pDO1}$ | | | | | | | | | | | | rep$_{pTEF3}$ | | | | rep$_{pS86}$ | | | |
| Efs 599 | 16 | USA | HH | 3 | | | | | rep$_{pKL0018}$ | | | | | | | | rep$_{pTEF3}$ | | | | rep$_{pS86}$ | | | |
| IBUN9046YE | 16 | COL | PF | 3 | rep$_{pDO1}$ | | | | | | | Δrep$_{pTEF1}$ | | | | | | | rep$_{pGB354}$ | | | | | |
| 712T | 476 | TUN | CF | 3 | rep$_{pDO1}$ | | | | | | | Δrep$_{pTEF1}$ | | | | | rep$_{pTEF3}$ | | | | | | | |
| 697T | 476 | TUN | RC | 3 | rep$_{pDO1}$ | | | | | | | Δrep$_{pTEF1}$ | | | | | rep$_{pTEF3}$ | | | | | | | |
| 18026 | 476 | CHI | HP | 3 | rep$_{pDO1}$ | | | | | | | Δrep$_{pTEF1}$ | rep$_{pTEF2}$ | | | | | | | | | | | |
| P.En250 | 476 | MAL | PF | 2 | | | | | | | | Δrep$_{pTEF1}$ | | | | | rep$_{pTEF3}$ | | | | | | | |
| 11340 | 476 | CHI | HP | 3 | | | | | | | | Δrep$_{pTEF1}$ | rep$_{pTEF2}$ | rep$_{pTW9}$ | | | | | | | | | | |
| 27149 | 858 | CHI | HP | 1 | | | | | | | | | | | | | | | | | | | | rep$_{pHY}$ |

Abbreviations: ST, sequence type; *n* , number of replication initiation genes; CHI, China; COL, Colombia; ESP, Spain; FRA, France; MAL, Malaysia; TUN, Tunisia; USA, United States of America; CF, chicken feces; HH, healthy human; HP, hospitalized patient; PF, pig feces; RC, retail chicken; RCA, retail cattle; RP, retail pig; WW, wastewaters.

[a]Isolates are ordered and separated according with their clsutering in the phylogenetic tree (Figure 1).

[b]The diversity of the plasmids present in each isolate is organized according the genes coding for replication initiator proteins (RIP, represented as rep genes). The prototype of each plasmid type is represented in each cell and they included: rep$_{pDO1}$ (GenBank acc. no. CP003584 - CDS12738); rep$_{pGB354}$ (GenBank acc. no. U83488 - *repI* ; rep$_{SAP093A}$ (GenBank acc. no. NC_013309 - SAP093A_001; rep$_{pKK627}$ (GenBank acc. no. NC_014156 - *repU* ); rep$_{pKL0018}$ (GenBank acc. no. AB290882 - *repB* ); rep$_{EF62pC}$ (GenBank acc. no. CP002494 - prgW); rep$_{pEJ97-1}$ (GenBank acc.no. AJ490170 - repA); rep$_{pTEF1}$ (GenBank acc. no NC_004669 - *repA1* ); rep$_{pTEF2}$ (GenBank acc. no. AE016831 - *repA2* ); rep$_{pTW9}$ (GenBank acc. no. AB563188 - *repA* ); rep$_{pCF10}$ (GenBank acc. no. AY855841 - *prgW* ); rep$_{pRE25}$ (GenBank acc. no. NC_008445 - *orf1* ); rep$_{pTEF3}$ (GenBank acc. no. - AE016832 - CDS16); rep$_{pTW9}$ (GenBank acc. no. AB563188 - repE); rep$_{pGB354}$ (GenBank acc. no. U83488 - repR); rep$_{pMBB1}$ (GenBank acc. no. U26268 - *repB* ); rep$_{pS86}$ (GenBank acc. no. AJ223161 - *repA* ); rep$_{pE1_13}$ (GenBank acc. no. CP018068 - BO233_15710); rep$_{pE394}$ (GenBank acc. no. KP399637 - *repB* ); rep$_{pHY}$ (GenBank acc. no. AB570326 - *repA* ).

*optrA* When possible to assess the location of *optrA* on plasmids, the correspondent *rep* genes are marked with a cell surronded by red.

Replication initiation genes are presented in different grey tones according with the highest homology they present with strain references retrieved in PlasmidFinder or in our in-house databa

Homology: 100% ≥99%<100% ≥97%<99% ≥95%<97% ≥90%<95% ≥80%<90%

**Table S6.** Nomenclature and gene mutations of *optrA* variants.

| Name CGE | Duplications CGE | Strain | GenBank | | | PMID | Bender *et al* | Morroni *et al* | 7 | 34 | 119 | 240 | 311 | 335 | 366 | 404 | 405 | 474 | 526 | 582 | 606 | 739 | 766 | 849 | 856 | 860 |
|---|---|---|---|---|---|---|---|---|---|---|---|---|---|---|---|---|---|---|---|---|---|---|---|---|---|---|
| ***optrA_1*** | | E349 | KP399637 | | 25977397 | | *optrA* $_{WT}$ | WT | A | A | G | G | T | C | C | A | T | C | T | A | C | G | G | C | C | T |
| ***optrA_2*** | | E016, 05K173 | KT862781 | KX620938 | 26903276 | 27645239 | 30236952 *optrA6* | EDM | G | - | - | - | - | - | - | - | - | - | G | - | - | - | - | T | - | - |
| ***optrA_3*** | | XY17 | KT862780 | | 26903276 | | 30236952 *optrA3* | | - | - | - | - | - | - | - | - | - | - | - | - | - | - | - | - | - | - |
| ***optrA_4*** | *optrA_6* | E419, 13F074 | KT862777 | KX620940 | 26903276 | 27645239 | | KD | - | - | - | - | - | A | - | - | - | - | G | - | - | - | - | - | - | - |
| ***optrA_5*** | | E147 | KT862783 | | 26903276 | | 30236952 *optrA4* | | - | - | - | - | - | - | - | - | - | - | - | - | - | - | - | - | - | - |
| ***optrA_7*** | | G20, 09L128 | KT862784 | KX620937 | 26903276 | 27645239 | | EDD | G | - | - | - | - | - | - | - | - | - | G | - | - | - | - | T | - | - |
| ***optrA_8*** | *optrA_9, optrA_10* | 10-2-2, 09P292 | KT862775 | KX620935 | 26903276 | 27645239 | 30236952 *optrA2* | DP | - | - | - | - | - | - | - | - | - | - | G | - | - | - | - | - | - | - |
| ***optrA_11*** | *optrA12* | E079, 11C219 | KT862782 | KX620933 | 26903276 | 27645239 | 30236952 *optrA7* | DD | - | - | - | - | - | - | - | - | - | - | G | - | - | - | - | - | - | - |
| ***optrA_13*** | | E35048 | KT892063 | | 26702919 | 27645239 | | LEYYWDVDASKELYNKQLEIG | G | T | - | A | - | - | A | G | G | T | G | T | T | - | - | - | - | - |
| ***optrA_14*** | | wo28-3 | KT601170 | | 26953332 | 27645239 | | EYDD | G | T | - | - | - | - | - | - | - | - | G | - | - | - | - | - | - | - |
| ***optrA_15*** | | 13G340 | KX620936 | | 27645239 | | | ED | G | - | - | - | - | - | - | - | - | - | G | - | - | - | - | T | - | - |
| ***optrA_16*** | | 11C233 | KX620941 | | 27645239 | | | KDP | - | - | - | - | - | A | - | - | - | - | G | - | - | - | - | - | - | - |
| ***optrA_17*** | | 13G339 | KX620934 | | 27645239 | | | DK | - | - | - | - | - | - | - | - | - | - | G | - | - | - | A | - | - | - |
| ***optrA_18*** | | 11L002 | KX620942 | | 27645239 | | | RDK | - | - | - | - | G | - | - | - | - | - | G | - | - | - | A | - | - | - |
| ***optrA_19*** | | 11R356 | KX620939 | | 27645239 | | 30236952 *optrA8* | EYDNDM | G | T | - | A | - | - | - | - | - | - | G | - | - | A | - | T | - | - |
| ***optrA_20*** | | 34E | New | | This study | | | - | G | - | A | A | - | - | A | G | - | T | - | T | T | - | - | - | A | A |
| ***optrA_21*** | | Efs 599 | New | | This study | | | - | - | - | A | - | - | - | - | - | - | - | - | - | - | - | - | - | - | - |
| ***optrA_22*** | | Iso4Bar | New | | This study | | | - | - | - | - | - | - | - | - | - | - | - | G | - | - | - | - | - | - | - |
| ***optrA_23*** | | Iso5Bar | New | | This study | | | - | G | - | - | - | - | - | - | - | - | - | - | - | - | - | - | - | - | - |
| ***optrA_24*** | | P.En250 | New | | This study | | | - | G | - | - | - | - | - | - | - | - | - | G | - | - | - | - | T | - | - |
| ***optrA_25*** | | N60443F | New | | This study | | | - | G | T | A | A | - | - | - | - | - | - | G | - | - | A | - | T | - | - |
| ***optrA_26*** | | N48037F | New | | This study | | | - | - | - | - | - | - | A | - | - | - | - | G | - | - | - | - | - | - | - |
| **Amino acid sequence** | | | | | | | | | **K3E** | **N12Y** | **G40D** | **V83V** | **I104R** | **T112K** | **N122K** | **Y135C** | **Y135W** | **Y158Y** | **D176Y** | **I194I** | **D202D** | **N247D** | **E256K** | **D283D** | **Q286K** | **I287K** |

**Position** (continued)

| Name CGE | 874 | 894 | 1035 | 1041 | 1049 | 1178 | 1184 | 1186 | 1197 | 1231 | 1441 | 1449 | 1464 | 1491 | 1500 | 1525 | 1530 |
|---|---|---|---|---|---|---|---|---|---|---|---|---|---|---|---|---|---|
| ***optrA_1*** | A | T | G | A | C | G | T | G | T | A | A | T | A | A | T | C | T |
| ***optrA_2*** | - | - | - | - | - | - | - | - | - | - | - | - | - | - | - | - | - |
| ***optrA_3*** | - | - | - | - | - | - | - | - | - | - | - | - | - | - | - | - | - |
| ***optrA_4*** | - | - | - | - | - | - | - | - | - | - | - | - | - | - | - | - | - |
| ***optrA_5*** | - | - | - | - | - | - | - | - | - | - | - | - | - | - | - | - | - |
| ***optrA_7*** | - | - | - | - | - | A | - | - | - | - | - | - | - | - | - | - | - |
| ***optrA_8*** | - | - | - | - | - | - | - | - | - | - | C | - | - | - | - | - | - |
| ***optrA_11*** | - | - | - | - | - | A | - | - | - | - | - | - | - | - | - | - | - |
| ***optrA_13*** | - | C | A | G | T | A | C | T | C | - | - | A | G | C | C | A | C |
| ***optrA_14*** | - | - | - | - | - | A | - | - | - | - | - | - | - | - | - | - | - |
| ***optrA_15*** | - | - | - | - | - | - | - | - | - | - | - | - | - | - | - | - | - |
| ***optrA_16*** | - | - | - | - | - | - | - | - | - | - | C | - | - | - | - | - | - |
| ***optrA_17*** | - | - | - | - | - | - | - | - | - | - | - | - | - | - | - | - | - |
| ***optrA_18*** | - | - | - | - | - | - | - | - | - | - | - | - | - | - | - | - | - |
| ***optrA_19*** | - | - | - | - | - | A | - | - | - | - | - | - | - | - | - | - | - |
| ***optrA_20*** | G | C | - | - | - | A | - | - | - | - | C | - | - | - | - | - | - |
| ***optrA_21*** | - | - | - | - | - | A | - | - | - | - | C | - | - | - | - | - | - |
| ***optrA_22*** | - | - | - | - | - | - | - | - | - | - | - | - | - | - | - | - | - |
| ***optrA_23*** | - | - | - | - | - | - | - | - | - | - | - | - | - | - | - | - | - |
| ***optrA_24*** | - | - | - | - | - | - | - | - | - | - | - | - | - | - | - | - | - |
| ***optrA_25*** | - | - | - | - | - | A | - | - | - | G | C | - | - | - | - | - | - |
| ***optrA_26*** | - | - | - | - | - | - | - | - | - | - | - | - | - | - | - | - | - |
| **Amino acid sequence** | **I292V** | **R298R** | **R345R** | **V347V** | **A345V** | **G393D** | **V395A** | **A396S** | **S399S** | **S410G** | **T481P** | **V483V** | **E488E** | **G497G** | **N500N** | **Q509K** | **G510G** |

| Name CGE | 1533 | 1587 | 1621 | 1626 | 1654 | 1662 | 1678 | 1686 | 1693 | 1737 | 1779 | 1833 | 1840 | 1866 | 1879 | 1899 | 1919 | 1948 |
|---|---|---|---|---|---|---|---|---|---|---|---|---|---|---|---|---|---|---|
| ***optrA_1*** | A | A | C | T | A | T | A | A | C | C | G | C | G | A | A | T | A | A |
| ***optrA_2*** | - | - | - | - | - | - | - | - | - | - | - | - | - | G | - | - | - | - |
| ***optrA_3*** | T | - | - | - | - | - | - | - | - | - | - | - | - | - | - | - | - | - |
| ***optrA_4*** | - | - | - | - | - | - | - | - | - | - | A | T | - | - | - | - | - | - |
| ***optrA_5*** | - | - | - | - | - | - | - | - | - | - | A | T | - | - | - | - | - | - |
| ***optrA_7*** | - | - | - | - | - | - | - | - | - | - | A | T | - | - | - | - | - | - |
| ***optrA_8*** | - | - | - | - | - | - | - | - | - | - | - | - | - | - | - | - | - | - |
| ***optrA_11*** | - | - | - | - | - | - | - | - | - | - | - | - | - | - | - | - | - | - |
| ***optrA_13*** | - | G | G | C | T | C | T | T | A | T | - | - | C | - | T | A | T | G |
| ***optrA_14*** | - | - | - | - | - | - | - | - | - | - | - | - | - | - | - | - | - | - |
| ***optrA_15*** | - | - | - | - | - | - | - | - | - | - | - | - | - | - | - | - | - | - |
| ***optrA_16*** | - | - | - | - | - | - | - | - | - | - | - | - | - | - | - | - | - | - |
| ***optrA_17*** | - | - | - | - | - | - | - | - | - | - | - | - | - | - | - | - | - | - |
| ***optrA_18*** | - | - | - | - | - | - | - | - | - | - | - | - | - | - | - | - | - | - |
| ***optrA_19*** | - | - | - | - | - | - | - | - | - | - | - | - | - | G | - | - | - | - |
| ***optrA_20*** | - | - | - | - | - | - | - | - | - | - | - | - | - | G | - | - | - | - |
| ***optrA_21*** | - | - | - | - | - | - | - | - | - | - | - | - | - | - | - | - | - | - |
| ***optrA_22*** | - | - | - | - | - | - | - | - | - | - | - | - | - | - | - | - | - | - |
| ***optrA_23*** | - | - | - | - | - | - | - | - | - | - | - | - | - | - | - | - | - | - |
| ***optrA_24*** | - | - | - | - | - | - | - | - | - | - | - | T | - | - | - | - | - | - |
| ***optrA_25*** | - | - | - | - | - | - | - | - | - | - | - | - | - | G | - | - | - | - |
| ***optrA_26*** | - | - | - | - | - | - | - | - | - | - | - | - | - | - | - | - | - | - |
| **Amino acid sequence** | **T511T** | **L529L** | **Q541E** | **S542S** | **M552L** | **S554S** | **N560Y** | **K562N** | **Q565K** | **S579S** | **K595K** | **I611I** | **E614Q** | **I622M** | **I627L** | **D633E** | **N640I** | **R650G** |

Non-synonimous mutations
